# The G-Protein-Coupled Receptor Kinase 2 Orchestrates Hair Follicle Homeostasis

**DOI:** 10.1101/2024.04.11.589052

**Authors:** Alejandro Asensio, Maria Sanz-Flores, Kif Liakath-Ali, Julia Palacios-García, Jesús M Paramio, Ramon García-Escudero, Federico Mayor, Catalina Ribas

## Abstract

Tightly regulated cell-cell and cell-niche intercommunications via intertwined signaling networks are involved in maintaining normal hair follicle (HF) homeostasis, cycling and cell fate determination. However, knowledge of specific mechanisms by which hair loss takes place under pathological situations is needed. Using a keratinocyte-specific knockout mouse model, we uncover that the G-protein-coupled receptor kinase 2 (GRK2) signaling node plays a key role in HF homeostasis. Epidermal GRK2 ablation causes alterations during anagen induction, giving rise to abnormal cyst-like structures. HF-linked cysts display aberrant growth and differentiation patterns as well as lineage infidelity, displaying features of abortive HFs unable to fully acquire canonical hallmarks. Cysts triggered by GRK2 deletion displace the dermal papilla away from the bulge and promote irreversible changes in HF stem cell architecture, leading to bulge destruction and hair loss. Our data provide unforeseen roles of GRK2 in epidermal physiology and uncover mechanisms linking dystrophic follicular cysts formation with hair loss, with potential connections to pathogenic processes operating in immune-mediated alopecias.

## Introduction

Hair follicles (HFs) constitute one of the main appendages of mammalian epidermis. They are composed of epithelial cells and have been widely used as a “mini-organ” model system for the study of adult stem cells and their differentiation processes taking place during the hair cycle [1, 2]. Hair cycle consists of three stages: growth (anagen), degeneration (catagen) and resting phase (telogen) [3]. Throughout lifetime, postnatal hair follicles undergo several cyclic rounds of destruction and regeneration, setting a system in which stem cells of resting HFs proliferate and differentiate giving rise to the hair shaft and the rest of anagen HF populations, thus making this system an ideal setting to study adult stem cell dynamics [4]. Tightly regulated spatiotemporal cell-cell and cell-niche interactions are responsible for maintaining hair homeostasis, whereas malfunctioning of any of these factors can lead to hair growth disorders such as alopecias [5].

Hair follicle stem cells (HFSC) reside in a specialized niche known as the bulge and are in charge of fueling hair growth during the first stages of the anagen phase [6, 7]. During telogen, the epithelial part of the HF includes slow-cycling bulge cells and the Hair Germ (HG) [8], a compartmentalized cluster of cells only visible during the resting phase which separates the bulge form the underlying specialized mesenchyme, the Dermal Papilla (DP) [9]. Epithelial-mesenchymal interactions are crucial during hair cycle [10–12] and the location of DP in close proximity to the HG or to the regressing HF enables a constant crosstalk between these structures, leading to the concept of DP as an orchestrator of the hair cycle, instructing the epithelial part of the HF to enter in either the regenerative or the destructive phase of the cycle [13, 14]. Several studies have pointed that bulge and HG cells not only differ in terms of their proximity to the DP, but are molecularly distinct entities, being HG cells biochemically and epigenetically primed to respond to anagen-inducing cues arising from the DP [15–17].

Cell identity within HFs is maintained by master transcription factors responsible for inducing HF commitment whilst blocking the expression of interfollicular epidermis (IFE) set of genes which are functional in other cell lineages present in the epidermis [18]. During telogen to anagen transition, complex cell signaling networks such as Shh, Notch, BMP or TGF-β [19–25] coordinately operate to induce tissue growth, regeneration and differentiation. For instance, Wnt/β-catenin pathway has a prominent role in inducing hair growth [26–29]. HFSC trigger hair production in response to Wnt signaling, and β-catenin stabilization plays a central role in the shift from TCF3/4 repressors, which are active in quiescent SCs, to Lef1, present in activated SC poised to regenerate the cycling components of the HF [30, 31]. Terminal differentiation during SC commitment takes place in a sequential manner, being the outer root sheath (ORS) and the companion layer (CL) the first to be specified, with other lineages such as matrix and inner root sheath (IRS) being generated at later stages of anagen progression [32].

A better understanding of how the different components of signaling networks crosstalk and are integrated in normal HF homeostasis, cycling and cell fate determination is needed to identify key regulatory hubs in these processes and their alterations in pathological conditions, leading to hair loss. In this regard, previous studies have reported the growth of cyst-like structures as a result of ablating signaling components relevant for HF biology [27, 28, 33–39]. However, the mechanisms linking cyst growth with a hair loss phenotype are not well understood.

G-protein-coupled receptor kinase 2 (GRK2) was first identified as a G-protein-coupled receptor (GPCR) modulator. Its canonical role is phosphorylation of activated GPCRs, leading to the recruitment of β-arrestins, and subsequent β-arrestin-dependent receptor desensitization, internalization and signaling. In addition, GRK2 exhibits a complex cell type and context-specific repertoire of non-GPCRs partners and substrates, acting as a signaling node involved in multiple cellular processes. Through such complex interactomes, GRK2 participates in key cellular functions, whereas altered levels of this protein may play a role in the maladaptive rewiring of signaling networks taking place in different disease contexts, such as cardiovascular, tumoral or metabolic disorders [40–42].

We have previously reported that GRK2 plays a key role in preserving epithelial cell identity in the context of head and neck squamous cell carcinoma (HNSCC), which arises from the epithelium lining the oral cavity, pharynx, and larynx. GRK2 expression is reduced in undifferentiated, high-grade human HNSCC tumors, whereas its downregulation in model human HNSCC cells is sufficient to trigger mesenchymal features and enhanced malignant potential in mice models, suggesting that GRK2 is very relevant to maintain and safeguard an epithelial phenotype [43]. However, the role of this protein in HF homeostasis has not been investigated.

In this study, we utilized mouse genetics to uncover the biological role of GRK2 in this setting. We reveal that keratinocyte-specific ablation of GRK2 leads to major changes in anagen induction, hair cycling and cell lineage determination in adult mice, leading to the formation of aberrant epidermal cyst structures. Interestingly, cysts triggered by GRK2 deletion displace dermal papilla (DP) away from the bulge and promote irreversible changes in HFSC niche architecture, eventually leading to bulge destruction and hair loss. Finally, we find that cysts are destroyed during aging by mechanisms reminiscent to those described in immune-mediated alopecias. Overall, our study unravels a central role of GRK2 in hair follicle epithelial cell biology.

## Results

### Epidermal GRK2 deletion triggers progressive hair loss with age and early changes in hair follicle morphology along with the development of cysts upon adult hair cycling

To analyze the potential role of GRK2 in hair follicle maintenance in vivo, we specifically deleted the GRK2 gene in epidermal and hair follicle cells by crossing homozygous GRK2 floxed mice (*GRK2* ^F/F^) with animals harboring Cre recombinase under the Keratin14 promoter (*K14-Cre*), which directs Cre expression to Keratin14-expressing stratified epithelial compartments. Genomic characterization (Fig.S1a), immunostaining of dorsal skin sections (Fig.S1b) and western blot analysis of isolated epidermal cells (Fig.S1c) confirmed GRK2 deletion in both hair follicles (HF) and interfollicular epidermis (IFE) in the *GRK2^F/F^;K14-Cre* mice epidermal KO animals (named GRK2 eKO from now on), whereas littermates lacking K14-Cre expression and thus retaining GRK2 floxed alleles (designated as control mice throughout the manuscript) displayed normal expression levels of GRK2.

Interestingly, we noted that although young (2 months) control and GRK2 eKO littermates were undistinguishable and displayed a normal hair phenotype, GRK2 eKO mice showed signs of hair loss with age, with an already sparser hair coat by the age of 8 months, and marked hair loss, with numerous hairless regions at 18 months (Fig.1a). Such age-related hair loss process was confirmed at the histological level (Fig.1b). Importantly, we observed that all GRK2 eKO animals exhibited abnormal epidermal cyst-like structures appearing in the dermis or linked to the lowest part of HF. These cysts were already present in 2-month-old mice (Fig.1b insert), before the onset of macroscopic hair loss, and their number increased significatively thereafter, peaking at 8 months (Fig.1b).

**Figure 1.**
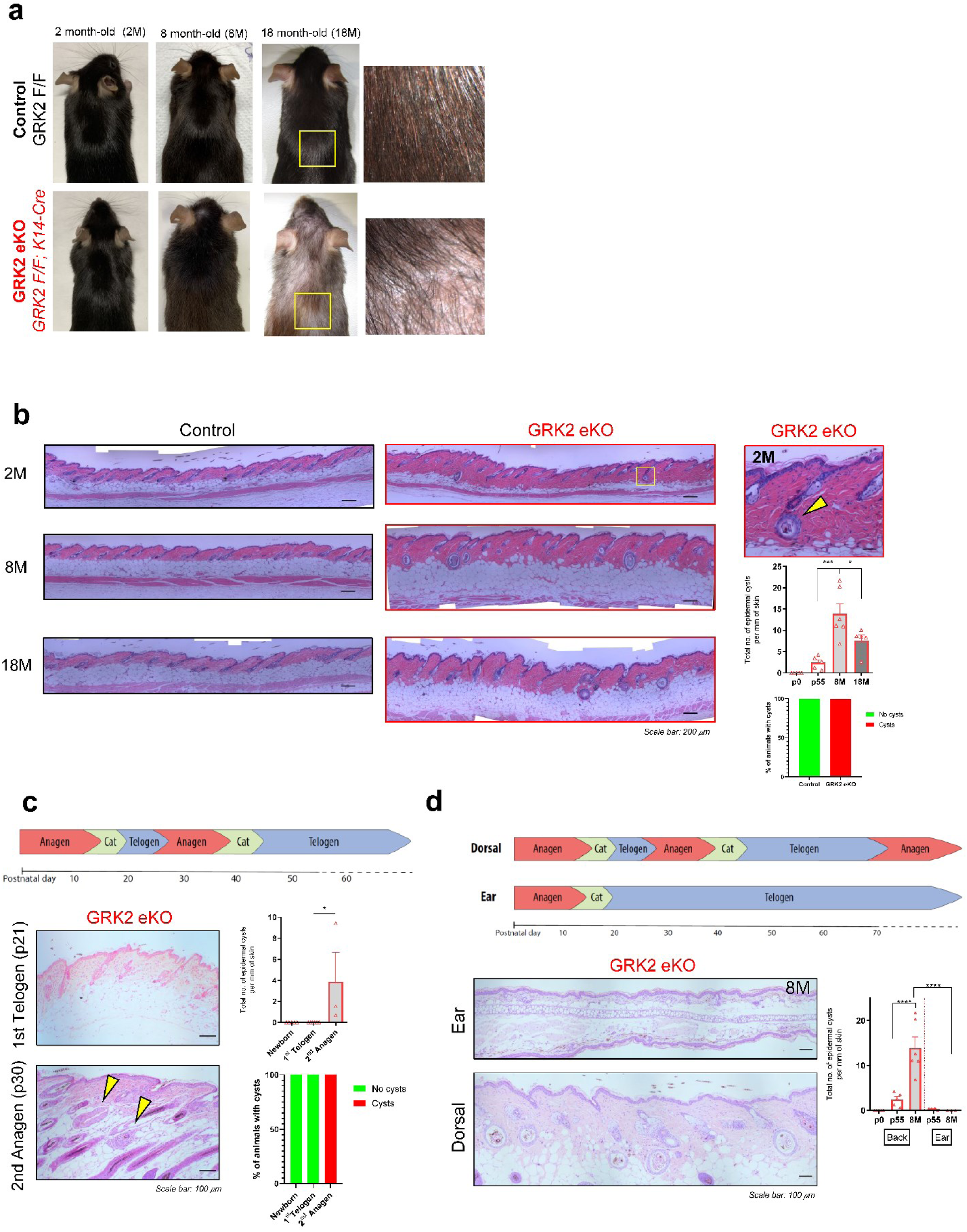
GRK2 eKO mice display progressive age-related hair loss with early changes in hair follicle morphology and cysts development related to hair cycling. (A) Comparative analysis of dorsal skin appearance at different ages, with detailed 18-mont-old view (right, square area amplified). (B) Representative skin sections showing cumulative alterations in hair follicle homeostasis including cysts at 2-month-old (square area amplified below) before macroscopic alterations are noted. Quantification of cysts/mm of skin observed in GRK2 eKO mice at different ages. As noted in the graph, all GRK2 eKO animals exhibited abnormal cyst-like structures at this stage. (C) Analysis of cyst presence at different ages showed that cysts are absent in newborn or 1st telogen mouse but show a penetrance of 100% after 2nd postnatal anagen. Quantification of cysts/mm of skin observed on GRK2 eKO skin revealed a significant increase in their numbers at p30 (2^nd^ anagen) compared to previous stages (Newborn and 1^st^ telogen). (D) Schematic view of the differential cycling properties of skin domains. The analysis of their growing pattern in GRK2 eKO animals revealed cyst numbers peak at 8-month-old mice and a preferential growth in cycling anatomical domains such as dorsal compared to resting ear skin. Means ± SEM data are represented, and statistical analysis was performed by 1-way ANOVA followed by Bonferroni’s post-hoc test, ****p< 0.0001.

To gain insights about the dynamics of cysts growth in GRK2 eKO animals, we examined skin sections at different stages of development. Importantly, GRK2 eKO pups are born without cystic structures, displaying an overall normal appearance of hair follicles both in length and number compared to controls (Fig.S2a). No cysts were observed either in newborns or at the first telogen stage at p21 (Fig.1c), when HF morphogenesis has been completed, pointing to a specific role for GRK2 during adult HF homeostasis and cycling. Consistent with this notion, all GRK2 eKO mice sections examined after second postnatal anagen (>p30) showed the presence of these abnormal structures (Fig.1c). As we pointed above, dorsal skin cyst numbers strongly increase by the age of 8-months in GRK2 eKO mice, when control littermates HF have usually undergone circa 4 cycles, and when macroscopic hair loss effects are apparent in the knockout animals (Fig.1d). Strikingly, cyst growth in adult GRK2 eKO animals was totally abrogated in ear skin, which remains in a refractory telogen-arrested state for extended periods of time [44] (Fig.1d). These data suggested that GRK2 may play a central role in adult HF cycling and homeostasis. To further support this notion, we used a conditional mouse model of global tamoxifen-inducible GRK2 ablation (*TxCre-GRK2^F/F^)*. Triggering GRK2 deletion at the stage of second telogen in 2-month-old mice lead with time to the same phenotype observed in our constitutive epidermal specific knockout model, with major presence of cyst structures and macroscopic hair loss (Fig.S2b), clearly supporting our hypothesis regarding the need of GRK2 during adult hair cycling.

### GRK2 is required for proper anagen induction and progression

We next investigated the impact of GRK2 deletion on different stages of the hair cycle. We took advantage of the synchronous second telogen status (resting phase) of mouse dorsal skin HF at p55, before macroscopic hair loss has started, and the possibility of triggering the synchronized anagen start via depilation.

During telogen, the epithelial part of the HF is composed of slow-cycling hair follicle stem cells (HFSC), that reside in a specialized niche known as the bulge, the secondary Hair Germ (HG) and a specialized mesenchyme, the Dermal Papilla (DP). We found that at p55 second telogen stage, both control and GRK2 eKO mice displayed non-significant differences in the amount of bulge HFSC, as assessed by either flow cytometry (CD34^+^ α6-integrin^+^) or immunofluorescence (CD34^+^, Sox9^+^ and Keratin 15^+^) (Fig.2a-b). Interestingly however, GRK2 eKO mice exhibited smaller HG and fewer P-cadherin^+^ HG cells per HF section (Fig.2b), suggestive of an altered activation pattern of this primed HFSC compartment.

**Figure 2.**
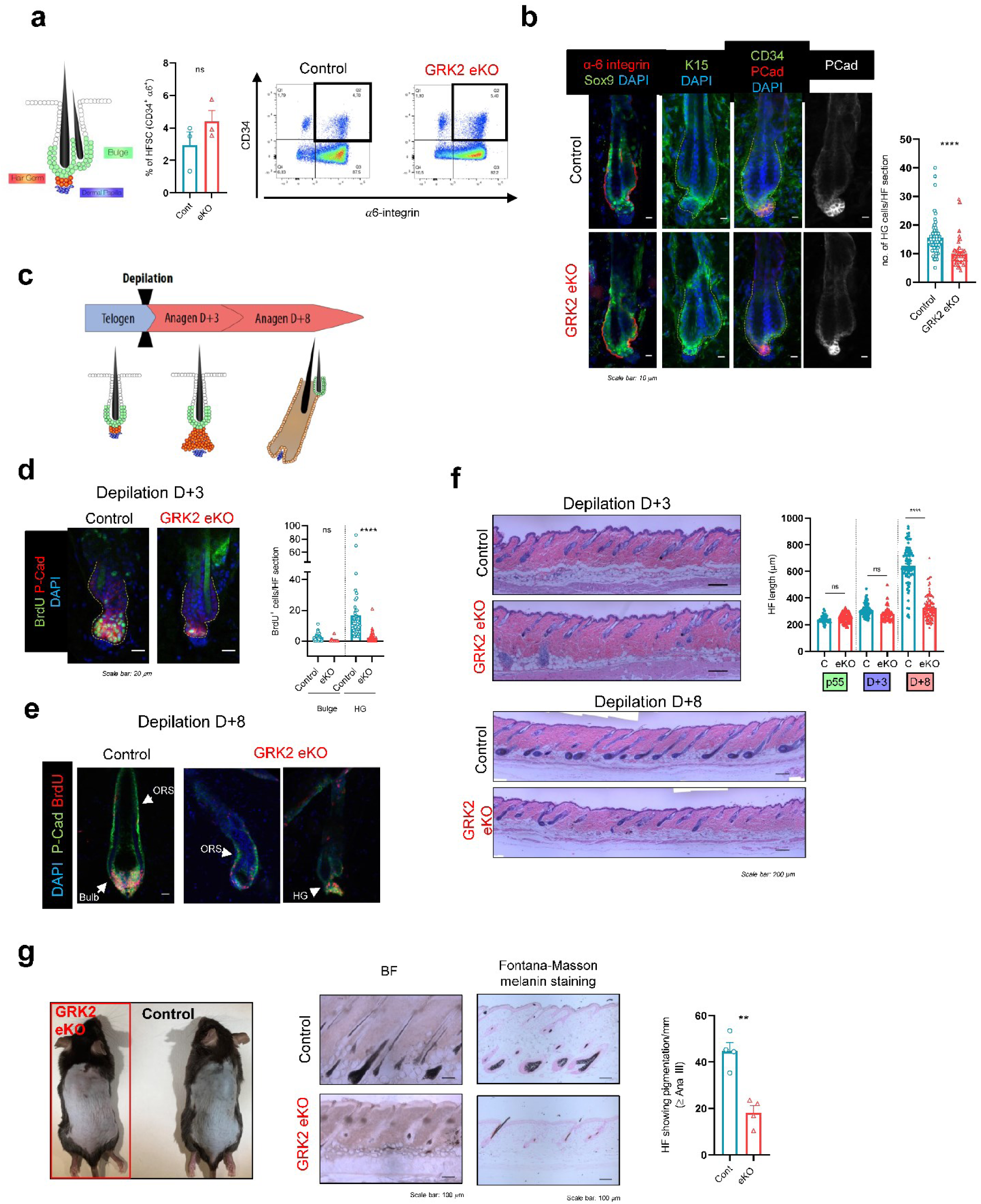
Young GRK2 eKO mice show conserved HFSC pool but deficient anagen induction. (A) Schematic representation of telogen HF structures. Second telogen GRK2 eKO mice show no changes in FACS-sorted CD34^+^ α6-integrin^+^ bulge cell numbers. (B) Immunofluorescence analysis confirmed unchanged numbers of bulge Keratin 15^+^, Sox9^+^ or CD34^+^ cells, in contrast to hair germs (HG), identified by P-cadherin, which are significantly smaller in GRK2 eKO mice. (C) Schematic representation of the depilation-induced hair growth experiment. Second telogen mice (p55) were shaved to induce synchronized growth of back skin HFs. Skin sections were analyzed at p55 and 3 or 8 days after hair removal. (D, E) Proliferation determined by BrdU incorporation is diminished in the HG of GRK2 eKO at early (D+3) (D) or later steps of anagen induction (D+8) (E) compared to control littermates. (F,G) Hair follicle length and presence of pigmented hair shafts were also significantly reduced in GRK2 eKO animals at D+8 stages. Representative images are shown. Means ± SEM data are represented, and statistical analysis was performed using unpaired t-test, **p < 0.01, **** p < 0.0001.

To explore the functional consequences of such alterations in the HG compartment, we depilated the p55 mouse back skin to induce the synchronized anagen onset and characterized HF status at 3 (D+3) or 8 (D+8) days post-depilation (Fig.2c). The initial regenerative phase of hair cycle implies proliferation of HG cells, eventually giving rise to the matrix by the anagen-III stage. Interestingly, proliferation of HG cells (BrdU incorporation) was clearly diminished in the absence of GRK2 at D+3 post depilation (Fig.2d), indicating a delay in the telogen-anagen transition. Such delay in anagen induction persisted at D+8 (Fig.2e). In control animals, the high levels of proliferation were confined to the hair bulb region, sustaining the formation of the hair shaft, and were indicative of canonical completion of anagen stages to a point when outer root sheath (ORS) cells have ceased their expansion (Fig.2e, left)). In sharp contrast, GRK2 eKO HFs displayed a retarded progression and were heterogeneous in their proliferation pattern and anagen status, ranging from early anagen-III HFs with emerging matrix and proliferative upper ORS (Fig.2e, middle) to anagen-I HFs where cell division can only be detected in a few HG cells (Fig.2e, right). Consistently, as judged by H&E staining, in contrast to the ≥Ana-III/IV HFs observed in control littermates 8 days post depilation, with a marked increase compared to D+3, HF length was significantly reduced in GRK2 eKO mice and remained unchanged compared to D+3 conditions (Fig.2f).

Pigmentation of mouse skin is tightly linked to the hair cycle stage and was also altered in GRK2 eKO mice. At 8 days post depilation, control and eKO skin displayed a different macroscopic appearance, with a darker grey coloration in control animals, indicative of advanced anagen stages (Fig.2g). Histological analysis and melanin staining quantification showed a significant decrease in the number of HFs displaying pigmentation in the region above the DP in GRK2 eKO mice (Fig.2g, right panels). Together, these data indicated that deletion of GRK2 in the epidermal compartment has a negative impact in anagen induction and hair cycle progression, pointing to postnatal anagen defects as one of the possible drivers of the progressive hair loss phenotype observed in GRK2 eKO mice.

HFs include a variety of cellular types with different roles throughout the cycle. To better understand the role of GRK2, we investigated the spatiotemporal dynamics of its expression in the HF. Second telogen single-cell transcriptomics (data obtained from published databases [45]) as well as our immunofluorescence staining (Fig.S3a) indicated that GRK2 is expressed in resting phase HF, with the highest signal found in the upper HF, upper bulge and HG. An additional study [16] also suggest a more dynamic expression of GRK2 transcript levels towards the late phases of telogen in HG compared to bulge cells (Fig.S3b). ATAC-Seq experiments [17] point to the same direction, with a peak of open chromatin identified in GRK2 genomic location in HG (Fig.S3c). At the protein level, we found that GRK2 levels increase over telogen bulge levels 3 days post depilation (Ana-I/II) in both bulge and HG of control HF. GRK2 levels persisted elevated in more advanced anagen stages (>Ana-III/IV), being significantly upregulated in all HF compartments and especially in the bulge, upper ORS and matrix (Fig.S3d). Altogether, it was tempting to suggest that physiological upregulation of GRK2 in HF is required for adequate anagen induction and progression, and that selective ablation in GRK2 eKO animals would trigger a delay in anagen onset and complex changes in hair cycling and HF homeostasis.

### GRK2 downregulation correlates with unbalanced status of key signaling pathways involved in anagen induction and hair cycling

Given the aforementioned induction of GRK2 expression, and the fact that GRK2 has been described as a central signaling hub [40], we hypothesized that the lack of this protein may alter the activation status of signaling pathways relevant in HF homeostasis and cycling.

Robust upregulation of the LEF1 transcription factor downstream the Wnt/βCatenin pathway takes place in the HG and its progeny during telogen-anagen transition, allowing a switch from the inhibitory TCF3/4 effectors in the bulge, opposed to β-Catenin signaling [31]. Of note, consistent with the delayed anagen observed p55 GRK2 eKO mice 3 days post-depilation, these animals displayed low levels of cytoplasmic β-Catenin as well as markedly reduced nuclear Lef1 levels in the HG region compared to controls (Fig.3a). These Wnt signaling reporters remained low even at 8 days post depilation, when some eKO HFs started developing the matrix.

**Figure 3.**
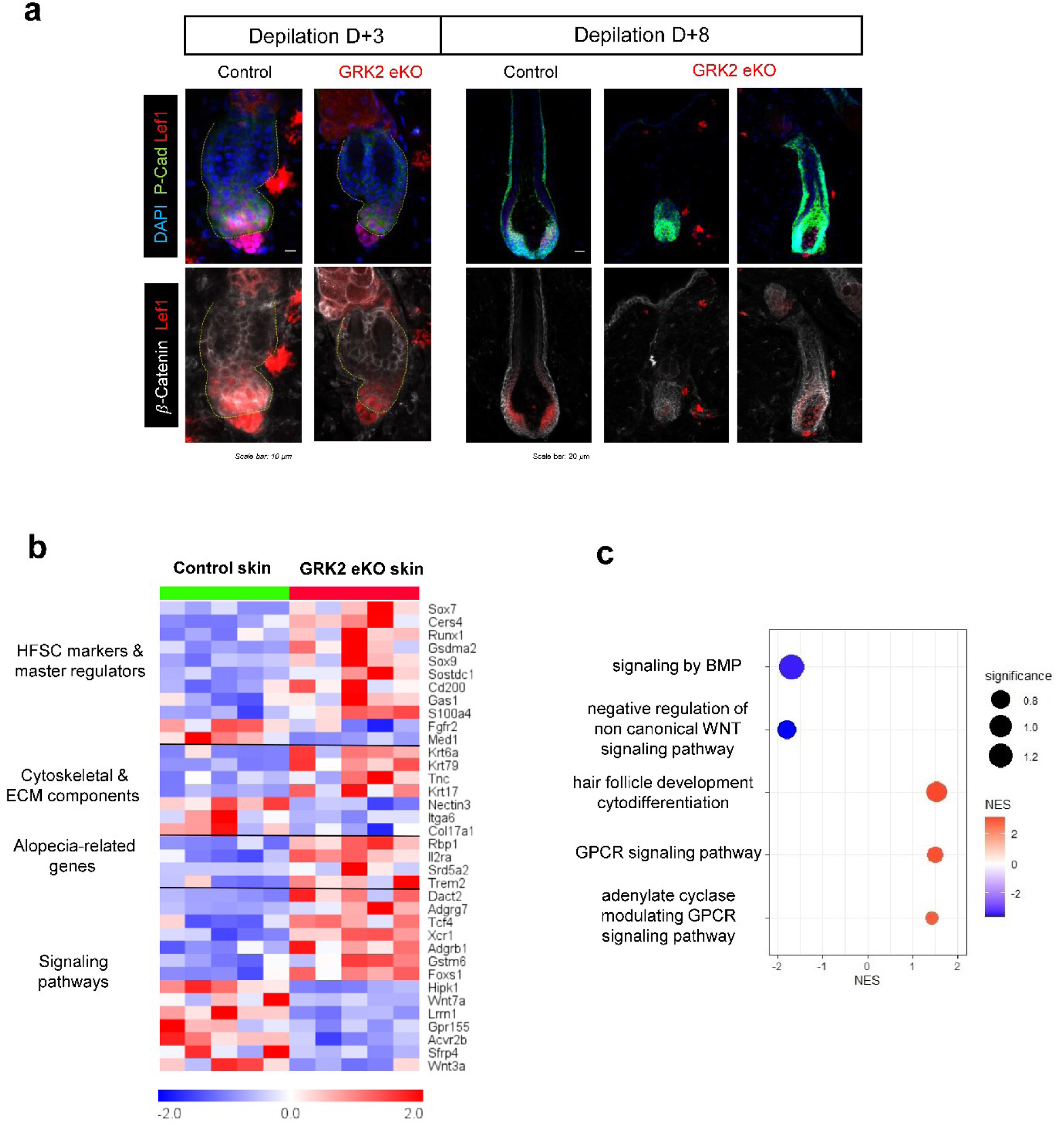
GRK2 downregulation leads to alterations in signaling pathways involved in stem cell maintenance. (A) Activation status of canonical Wnt/βCat readouts (βCatenin and Lef1) measured by immunofluorescence in p55 mice at different days post-depilation. A deficient activation of this pathway in GRK2 eKO anagen HFs at both D+3 or D+8 is noted compared to control mice. Representative images are shown. (B) Heatmap of differentially expressed genes between GRK2 eKO and Control skin from RNA-seq experiments. Differential expression was assessed from 5 skin samples in each group and upon DESeq2 analysis. (C) Graphical representation of GSEA analysis of RNAseq data from GRK2 eKO versus Control skin. Significant enrichments were observed in either Control (blue) or GRK2 eKO (red) for genesets related to hair follicle function, GPCR, or developmental signaling pathways. The significance is expressed as −log10(FDR).

To analyze the long-term outcome of the mentioned altered anagen process we performed an RNASeq experiment in 8-month-old mice whole skin to detect changes in transcriptional readouts of Wnt/βCatenin and other signaling pathways. Of note, several transcripts related to HFSC were both up or downregulated in GRK2 eKO animals compared to littermates, such as Runx1, Sostdc1, Gas1, Sox9 or Fgfr2 (Fig.3b). We also detected altered levels of relevant structural proteins regarding the cytoskeleton or extracellular matrix (ECM) such as Krt79, Tnc, Itga6 or Col17a1, which are involved in HFSC aging. It is noteworthy that BMP, Wnt, GPCR and adenylate cyclase pathway signaling components and signatures were either significantly enriched or decreased between genotypes (Fig.3c). The forementioned imbalance in signaling pathways involved in HFSC maintenance was exemplified by the significant decrease in key transcripts such as Wnt3a, Wnt7a or Acvr2b, or the increase in Tcf4 in GRK2 eKO skin (Fig.3b), overall translating into flawed “hair follicle development cytodifferentiation” signature (Fig.3c) and consistent with the alterations noted above in genes related to HFSC (Fig.3b) and to the GSEA analysis (Fig.S4). Overall, it was tempting to suggest that such alterations in cell signaling components may cooperatively act to trigger hair cycling defects and cyst emergence observed in this mouse model.

### Cysts are uncoupled from normal hair follicle differentiation and cycling features

A key feature of the GRK2 eKO phenotype was the presence of abnormal cyst-like structures appearing in the dermis or linked to HF in adult mice. Since GRK2 downregulation leads to deficient anagen induction and aberrant HF cycling, we reasoned that cysts may arise in constantly cycling mouse back skin because of the accretion of such altered features over time. We thus investigated the expression of key molecular markers of HF layers in the cyst’s structures (see scheme of cell layer markers found in an anagen phase hair follicle, Fig. 4a).

**Figure 4.**
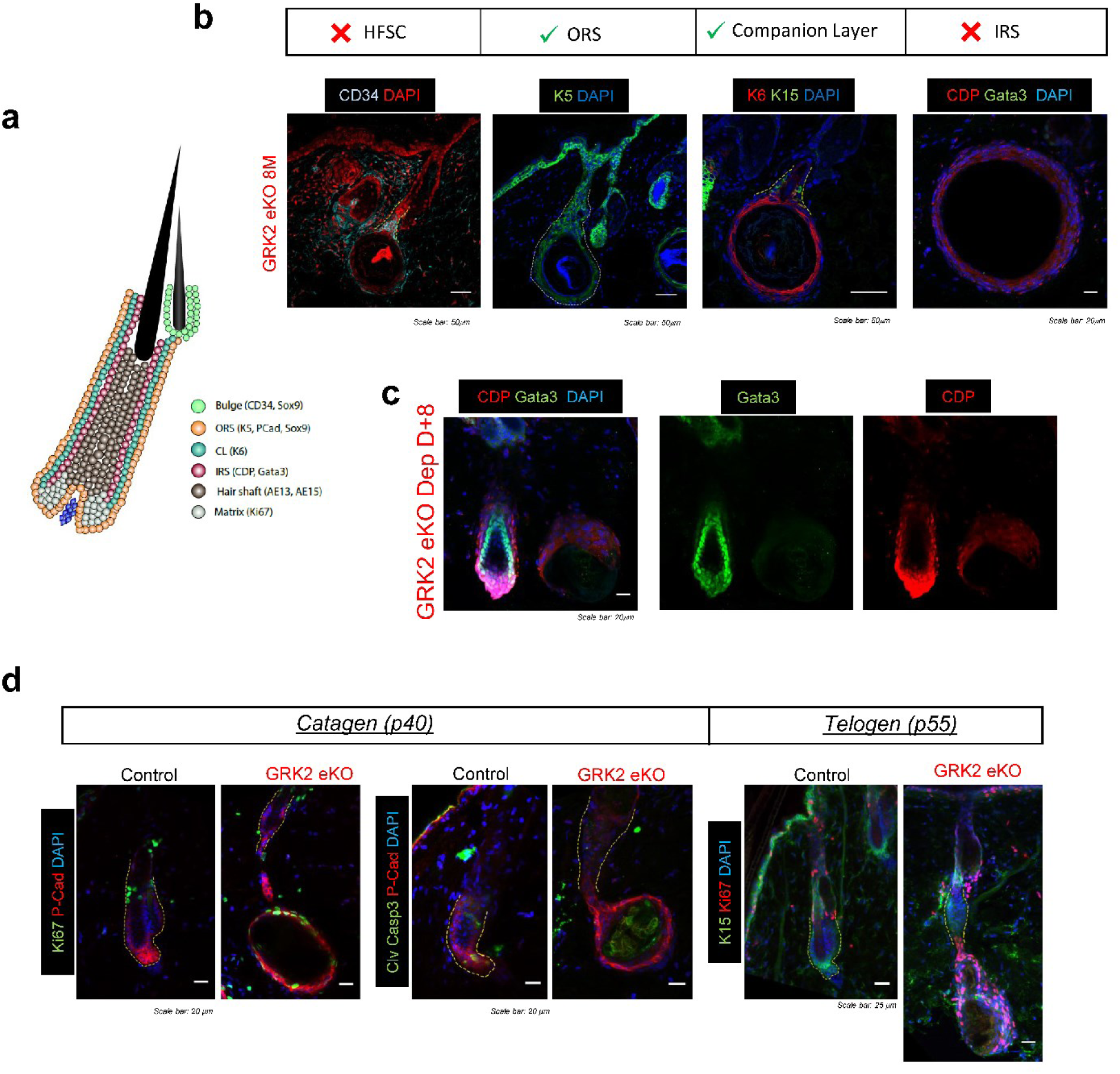
GRK2 eKO cysts arise from HFSC but are uncoupled from normal hair follicle differentiation and cycling features. (A) Representative schematic of concentric cell layers and their molecular markers thar are normally found in an anagen phase hair follicle (HF). (B) Immunofluorescence analysis of GRK2 KO skin sections reveals that cysts have an epidermal origin (Keratin 5 positive) and appear linked to the stem cell niche (CD34 positive). These structures form a continuous Keratin 6 positive layer with the inner bulge cell niche but do not express the classic HFSC markers CD34 or Keratin 15. Cyst appear as Gata3/CDP negative, indicating the lack of IRS cell fate. (C) Immunodetection of IRS and IRS/Matrix markers Gata3 and CDP reveal a lack of full anagen HF differentiation process in HF-derived cysts (right) compared to normal HF in the same GRK2 eKO animal shown in the left part of the image. (D) During synchronized hair cycle stages, unlike control littermates, GRK2 eKO cysts are proliferative (Ki67^+^) in catagen and bypass hair cycle regression remaining negative for apoptosis markers (caspase 3) during regression phase. During telogen, cysts are also Ki67^+^, which indicates that proliferation within these structures is not arrested during the resting phase. Representative immunofluorescence images are shown.

When skin sections were stained for HFSC markers, we observed that cysts appeared to grow from the lowest part of the HF (HG) and remained attached to the CD34^+^ bulge, although cysts structures displayed no labeling for this HFSC marker (Fig.4b). The epidermal origin of cysts was indicated by their positive staining for Keratin 5 and high Keratin 6 labeling, whereas the lack of Keratin 15 and dim α6-integrin staining confirmed their differential features from HFSC clusters (Fig.4b and Fig.S5a). The K6^+^ cell layer formed a continuous structure from the inner bulge stem cell niche, extending to the whole cyst (Fig.4b). Regarding anagen HF differentiation patterns, while ORS markers as K5 or P-cadherin were present in the cysts, as was the matrix ascending companion layer marker K6, the labeling of IRS determinants was markedly decreased (GATA3) or showed a non-nuclear unspecific distribution (CDP) in GRK2 eKO HF-linked cysts compared to normal HF from the same animal (Fig.4b-c). Of note, cysts showed on the inside an accumulation of eosinophilic substances that are likely keratin debris reminiscent of hair shafts and also contain lipid secretions marked by the Nile Red lipid dye (Fig.S5b), suggesting that these anomalous structures arise from HFSC clusters and remain attached to the niche yet displaying an aberrant, non-canonical differentiation program. Moreover, cyst walls appeared as a multilayered epithelium, occasionally composed by cells with enlarged and atypic nuclear morphology. (Fig.S5c). Overall, these features suggested that cysts showed features of abortive HFs unable to fully acquire the canonical anagen cell layers.

Strikingly, we observed that HF-derived cysts are not destroyed in catagen and proliferate steadily in a hair cycle stage-independent way. At p40, control HFs are synchronized in the regression phase (catagen), when no active proliferation (no Ki67) but rather destruction (caspase3) of the transient SC progeny generated during the anagen growth phase takes place (Fig.4d). However, at this catagen stage GRK2 eKO cysts cells are proliferating and contain no Caspase3^+^ apoptotic cells, suggesting that cyst cells are unresponsive to catagen-inducing cues, thus bypassing the regression phase. Furthermore, in the p55 telogen resting phase, contrary to control HFs, active proliferation was detected in cysts (Fig.4d). Overall, these data indicated that despite being structures originated from HFs and in many cases remaining linked with them, cyst cells display an altered fate, showing growth and differentiation patterns uncoupled from normal hair cycle features.

### GRK2 eKO cysts display lineage infidelity and tumoral-like features

Epidermis is comprised of two main cell lineages, the interfollicular epidermis (IFE) and HF, with differential features and cycling properties. Opposed to the cyclic and niche-induced proliferative rounds of HF cells, interfollicular SC undergo continuous proliferation to replenish cells lost in the skin surface through desquamation. Since the proliferation patterns of cysts were reminiscent to that of IFE, we investigated the expression of key epidermal lineage determinants in our aberrant structures in 8-month-old mice. Klf5 and Sox9 are key transcription factors (TFs) demarcating IFE and HF lineages, respectively, and under homeostatic conditions its presence appears to be mutually exclusive in the same cell [18]. These TFs govern cell identity by globally controlling lineage-specific expression profiles. Strikingly, cysts showed marked staining for the IFE Klf5 TF, while remaining positive for Sox9, indicative of their HF origin. In contrast, control animals or GRK2 eKO bulges showed expression of Sox9 alone (Fig.5a and Fig.S6a). The notion of an IFE-like phenotype and lineage infidelity in cysts was further strengthened by the expression of the IFE ΔNp63 marker (Fig.5b), and an active, IFE-like proliferation pattern as assessed by BrdU incorporation (Fig.S6b). Also indicative of the altered fate of cysts, cells were positive for the infundibulum marker Lrig1, expressed only in the upper HF in control animals (Fig.5c), whereas they were negative for junctional zone markers such as Gata6 (Fig.S6c). Of note, despite displaying expression of several IFE-specific TFs and being multilayered structures, the epidermal differentiation program was not fully completed in the cysts, that lacked key markers such as K10, Filaggrin or Loricrin (Fig.S6d). This probably reflects the opposing roles and fates triggered by the HF and IFE determinants co-expressed within the cysts.

**Figure 5.**
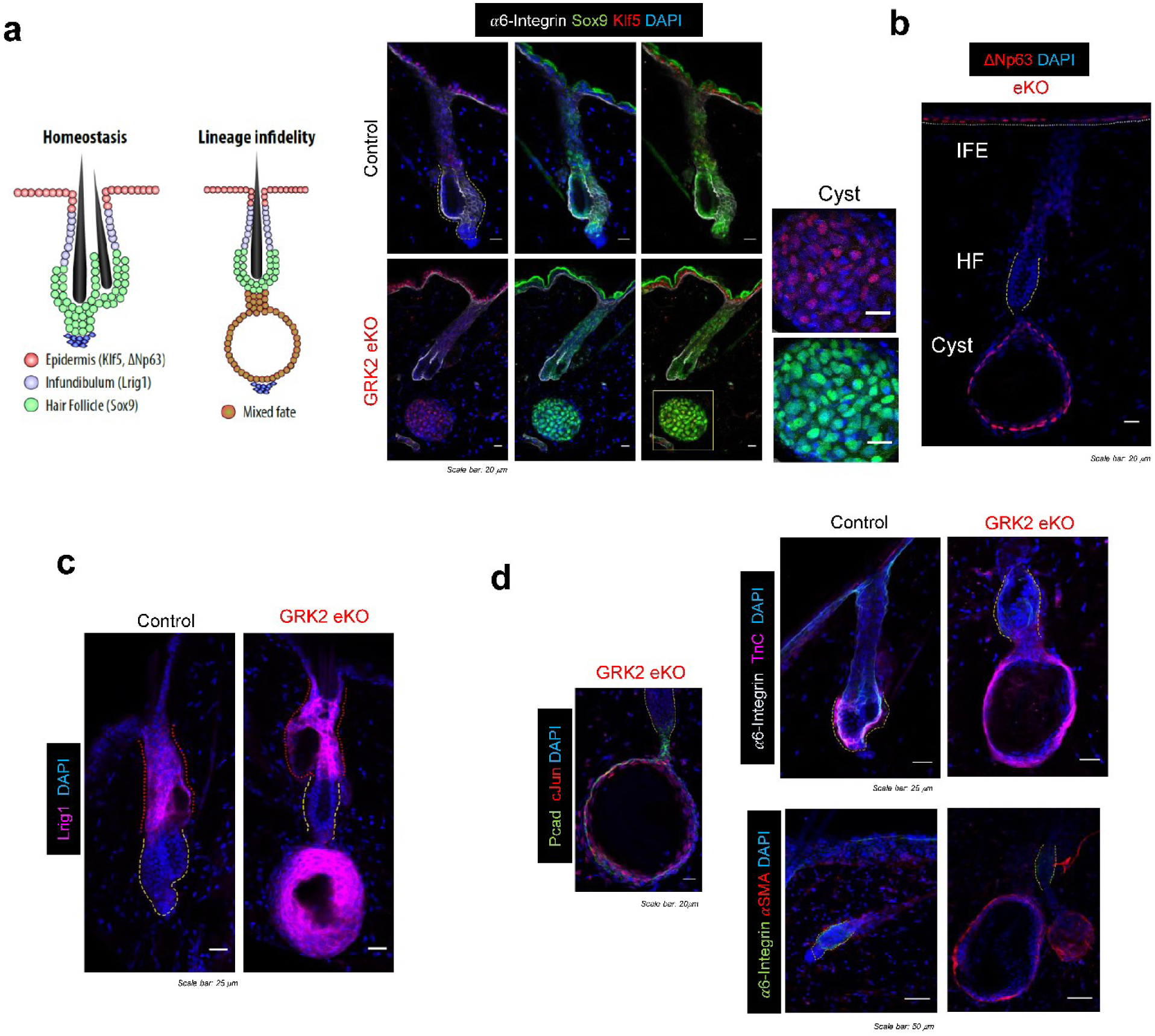
Cysts display lineage infidelity, co-expressing a mixture of IFE, HF and SCC-related tumoral markers. (A) Left, schematic representation of epidermal lineage markers. Right, immunofluorescence analysis shows the presence of cysts cells co-expressing Klf5 and Sox9 transcription factors (detailed in the inset), otherwise distinctly expressed in IFE and HF control cells, suggesting a lineage infidelity phenotype in these cells. The remaining Sox9 expression displayed by cysts cells further supports their follicular origin. (B) Mixed IFE and HF phenotype in cysts is further suggested by the presence of ΔNp63, an IFE marker. (D) Lineage infidelity features in GRK2 eKO cysts are shown together with the expression of tumor-associated markers, as the transcription factor c-Jun or extracellular matrix components such as TnC or αSMA, typical of pro-tumoral environments.

Co-expression of Klf5 and Sox9 has been described transiently during wound resolution or on a permanent basis in tumoral conditions in squamous cell carcinomas (SCCs) [18, 46]. Given that in the cysts appearing in GRK2 eKO mice, lineage infidelity takes place in “basal” conditions and is persistent over time, we searched for the presence of tumoral-related markers. Interestingly, cysts were markedly positive for the stress TF c-Jun (Fig.5d), which might contribute to the convergent expression of Sox9 and Klf5 [18]. In addition, cysts were positive for matrix remodeling or differential matrix deposition markers related to tumoral microenvironment hallmarks as TenascinC or αSMA (Fig.5d). However, despite displaying several tumoral features, GRK2 eKO mice remained skin tumor-free up to 20-month-old (Fig.S6e).

Altogether, our results put forward a crucial role of GRK2 in HF homeostasis and cycling, since its absence promotes a variety of alterations, ranging from deficient anagen induction to abnormal cell fate determination, eventually giving rise to the accumulation of aberrant HF-derived cysts uncoupled from canonical HF behavior. These data raised the questions of how cyst growth promotes progressive hair loss in GRK2 eKO mice and why cysts do not end up developing tumors despite presenting some tumoral features.

### Cyst growth displaces dermal papilla (DP) away from the bulge and irreversibly affects HFSC niche architecture leading to hair loss

The mesenchymal structure located below the HFSC niche known as Dermal Papilla (DP) has been reported to be crucial in orchestrating transitions between hair cycle stages, and its proximity to the HG at the proximal part of the telogen HF enables epithelial-mesenchymal interactions [47]. During normal hair cycle, anagen HF surrounds and “pushes” the DP downwards followed by a catagen phase that returns DP to its natural place, below the bulge in telogen (Fig.6a). In sharp contrast, in GRK2 eKO mice the inability of cysts to respond to catagen-inducing signals kept the DP in contact with the cysts but away from the HFSC niche in both HF-linked cysts or in already detached structures (Fig.6b). Since the telogen HF relies in a close association between DP and HG to efficiently induce the entry into the growing phase, the fact that cysts grow out from the HG region and displaces the DP would lead cyst-bearing HF to become irreversibly telogen-arrested.

**Figure 6.**
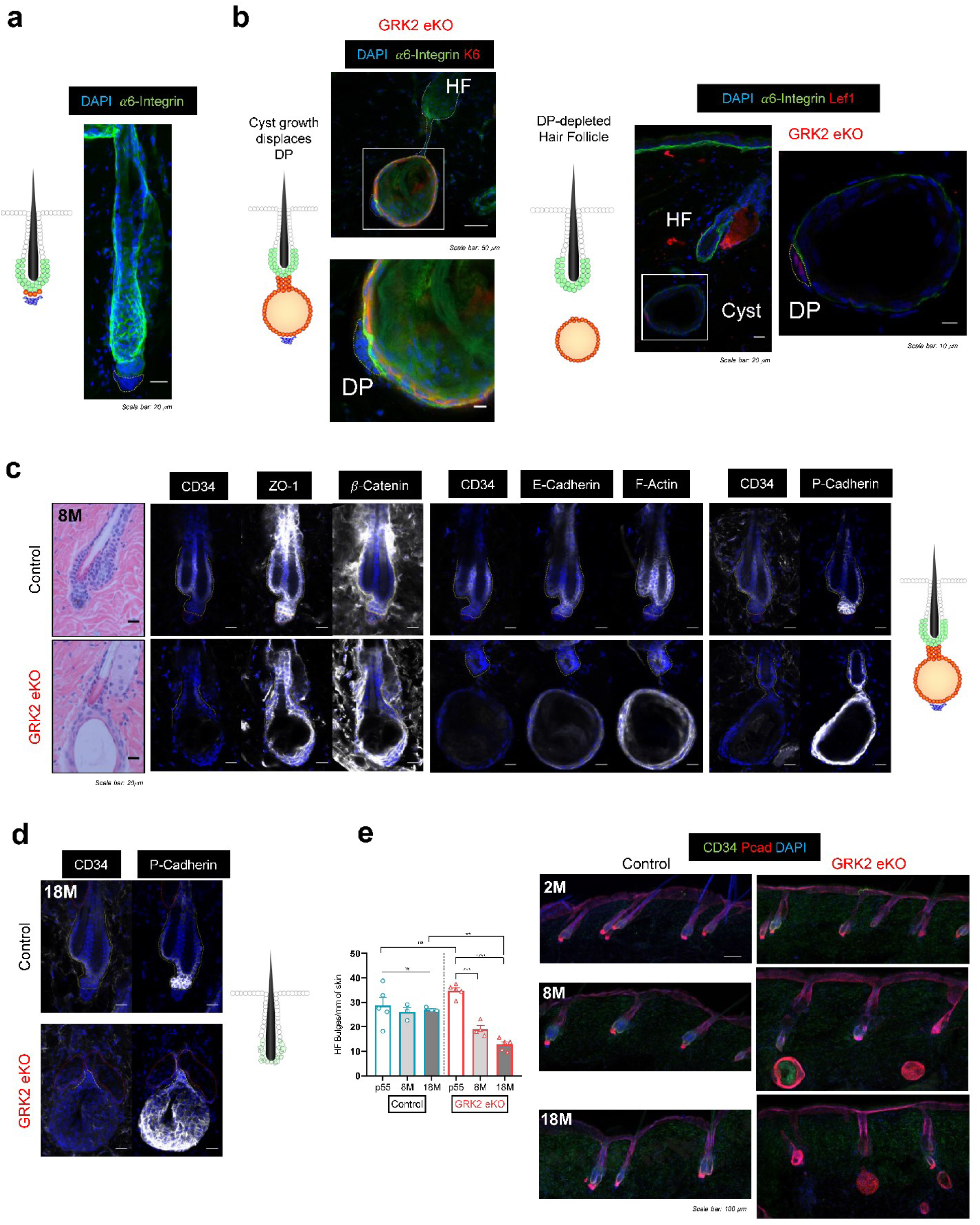
Dermal Papilla displacement and HFSC structural abnormalities couple cyst emergence with hair loss. (A) Normal HF morphology with the DP separated from the epidermal compartment by the α6-integrin^+^ basement membrane. (B) Histological analysis of 8-month-old GRK2 eKO mouse skin with the indicated markers showed that cyst growth from the lowest part of the HF displaces the DP (identified as Lef1^+^) away from the stem cell niche. Representative immunofluorescence images are shown. (C) Histochemical analysis of 8-month-old skin sections with the indicated markers showed miss-organization of both bulge cell layers which are no longer visible and appear interspersed in cyst-linked GRK2 eKO HFs. Marked structural alterations in bulge structure at 8-month-old GRK2 e KO mice were detected with immunofluorescence staining of key cell-cell junction proteins and cytoskeletal components. (D) These alterations led to loss of CD34 HFSC canonical marker in 18-month-old mutant mice compared to controls, together with the disappearance of the classical bulge morphology. (E) Quantification of bulge numbers revealed a decline in their amount specifically in GRK2 eKO animals during aging, a phenomenon not observed in control littermates. Representative immunofluorescence images are shown. Means ± SEM data are represented, and statistical analysis was performed by 1-way ANOVA followed by Bonferroni’s post-hoc test, **p < 0.01, ***p < 0.001, ****p < 0.0001.

As hair cycles progress during the lifetime of the mouse, HF have the potential to add one new bulge every cycle [48], so during the first telogen HFs would be single bulged and in second telogen (p55) can display two bulges. Such unique mechanism in mouse HF, termed club hair retention, enables hair coat maintenance. Interestingly, we observed in 8-month-old GRK2 eKO mice the presence of triple-bulged HF linked to cysts, had undergone at least three hair cycles before giving rise to a cyst and the subsequent depletion of DP, indicating that these GRK2 eKO HFs had undergone at least three hair cycles before giving rise to a cyst and the subsequent depletion of their DP (Fig.S7a). Of note, cysts linked to multi-bulge HFs were always detected to be in contact with the newest bulge (Fig.S7b), consistent with the notion of irreversible telogen arrest and hair growth impairment following cyst-induced DP displacement in GRK2 eKO animals.

We next explored the long-term consequences in bulge homeostasis and architecture triggered by aberrant cyst growth. Organization of HFSC within the niche is crucial to maintain SC function and genes related to ECM and bulge structure have been suggested to be critical during aging [49]. Control bulges display the typical bulge architecture comprised of two cell layers, the outer one essential during the regeneration process and the inner layer acting mainly as an anchoring point of the club hair to the HF and promoting quiescence through anagen inhibitory signals [4]. In contrast, histological examination of 8-month-old GRK2 eKO HFs giving rise to cysts showed a disorganized pattern, where the two bulge cell layers were not well distinguished and a pale and more eosinophilic labeling points to an altered differentiation state (Fig.6c). Co-immunofluorescence staining confirmed that cyst-linked HFs show structural alterations in the bulge region correlating with the loss of canonical HFSC markers. In control HF bulges, the outer cell layer was enriched in both P- and E-cadherins, whereas inner bulge cells only expressed E-cadherin. In contrast, cyst-linked GRK2 eKO HFs displayed a continuous and delocalized staining for both cadherins, coherent with the observed miss-organization of the outer and inner bulge cell layers that seem to be interspersed together. Consequently, adherens junctions, which rely on cadherin-cadherin homophilic interactions, were aberrantly distributed in GRK2 eKO compared to controls, as judged by ZO-1 and β-Catenin staining. Moreover, the inner bulge enrichment in F-actin staining detected in control animals was lost in GRK2 eKO littermates, indicating that alterations also extended to the cytoskeletal architecture and reflecting that structural miss-organization affects the whole bulge. Accordingly, bulges affected by these architectural changes triggered by cyst outgrowth in GRK2 eKO mice had lost most of CD34 expression compared to controls (Fig.6c). The analysis of aged (18-month-old) mouse skin sections further indicated that HFs linked to cysts were not only CD34 negative, but had progressed further in the degeneration process, displaying no obvious bulge structure in the lower region of the HF (Fig.6d), thereby triggering a striking decline in GRK2 eKO bulge numbers throughout aging as hair cycles take place (Fig.6e).

Consistent with the overall notion of delayed HF cycling, cyst formation in a fraction of HFs, altered niche architecture and hair loss in GRK2 eKO mice, the analysis of GRK2 knockout mice until day 20 post-depilation at p55 revealed that anagen induction resulted in a notable increase in cyst numbers when compared to that of 2^nd^ telogen phase (Fig.S7c). In this line, we noted that repetitive depilation, that would trigger several rounds of hair growth, exacerbates the GRK2 eKO phenotype. GRK2 eKO mice did grow a sparser hair coat than control animals when subjected to repetitive depilation. In contrast to control littermates, hair coat recovery was already severely compromised after 6 rounds of depilation in GRK2 eKO (Fig.S7d), and these depilated mice showed marked hair loss and a patent decrease in both HFSC CD34^+^ bulge cells and in total bulge structures by the end of the experiment after 9 depilation rounds (Fig.S7e). These data indicated that repetitive depilation aggravates stem cell exhaustion in GRK2 eKO mice correlating with the inability to regrow hair, and mechanistically argues in favor of the proposed model of HFSC niche architecture alterations and bulge destruction as a driver of hair loss in these animals.

Overall, our data indicate that cysts have long-term consequences on the HFSC niche architecture in GRK2 eKO mice, eventually leading to loss of bulge homeostasis and hair loss. According to our hypothesis, the more hair cycles that occur as mice age, the more HFs would be converted to cysts, thereby compromising DP-bulge interactions and bulge functionality, leading to the phenotype observed in GRK2 eKO mice.

### Epidermal cysts are destroyed during aging by mechanisms resembling immune-mediated alopecia

Our data thus far pointed to a very relevant role of GRK2 in HF homeostasis, since its absence leads to deficient anagen induction, abnormal cell fate determination, and accumulation of aberrant HF-derived cysts uncoupled from canonical HF behavior, eventually promoting structural changes in the HFSC niche, bulge destruction and hair loss.

Intriguingly, the persistence of lineage infidelity in basal conditions within cysts was reminiscent of that observed during wound healing or epithelial tumorigenesis [18]. However, cysts did not end up developing tumors throughout aging in GRK2 eKO mice, despite the presence of several tumoral markers, suggesting the existence of compensatory or counteracting processes. In fact, we observed that the number of cysts in GRK2 eKO animals reached a peak at 8 months of age and declined in 18-month-old mice, suggesting that HF-derived cysts are destroyed during aging (Fig.7).

**Figure 7.**
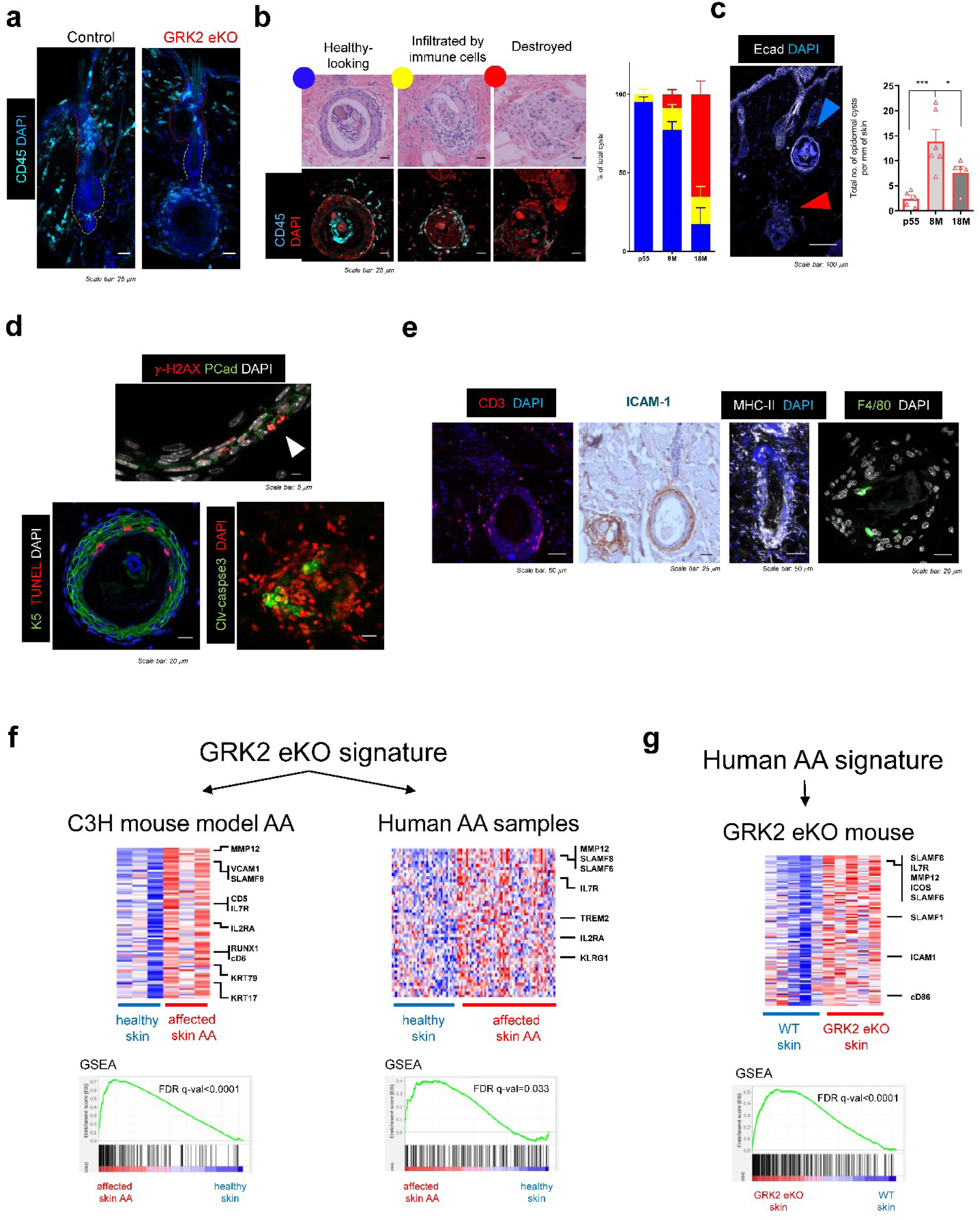
Epidermal cysts are destroyed during aging in a mechanism resembling immune-mediated alopecia. (A) Right graph shows that total number of cysts peak at 8M of age and then decrease by 18M. Left image indicates that cysts but not bulges attract immune infiltrate. In contrast to controls, where the presence of immune cells is restricted to the infundibular region, cysts are largely infiltrated by CD45^+^ immune cells. (B) According to their morphology and structural integrity, cysts were classified in “healthy” ones (blue) with round morphology and surrounding CD45^+^ cells, “compromised” cysts (yellow) that have started losing their morphology and are infiltrated by immune cells, and “destroyed” ones (red) formed by remnants free in the dermis. The number of the two later types increases with age, consistent with the evolution of the number of these structures with age shown in panel above. (C) Destroyed cysts (red arrow in lower right panel) are observed as structures lacking intercellular E-cadherin junctions, in contrast to the healthy-looking ones (blue arrow) that are positive for this marker. Quantification of cysts/mm of skin observed in GRK2 eKO mice at different ages. (D) Apoptotic cell markers such as cleaved caspase 3 or TUNEL were detected together with *γ*-H2AX within destroyed cysts, indicating that they are destroyed during aging. (E) Immune cells infiltrating cysts assessed with the indicated specific markers. CD3^+^ T cells are located in the surrounding dermis, consistent with the observed high expression of T-cell adhesion molecules (ICAM1). Antigen Presenting Cells (MHC-II^+^) and F4/80^+^ macrophages were also detected in infiltrated cysts. ((F) A gene expression signature of overexpressed genes in GRK2 eKO (GRK2 eKO signature) was enriched in alopecia samples obtained in the C3H mouse model (left panel), and in skin punch biopsy samples from AA patients (right panel). (G) A gene expression signature of overexpressed genes in human AA patients (human AA signature) was enriched in GRK2 eKO skin. Box and whiskers plots are used with means and min to max data are represented, and statistical analysis was performed by 1-way ANOVA followed by Bonferroni’s post-hoc test, ** p < 0.01, *** p < 0.001.

In the context of epithelial cell tumorigenesis, immune-epithelial interactions are of key importance [50]. In control mice, the presence of immune cells was restricted to the infundibular region, whilst the bulge displays what is termed the immune privilege [51]. However, in mutant animals CD45-positive immune cells were not only located in such area, but largely present in the vicinity of HFs and cysts, either inside cysts or surrounding them (Fig.7a-b).

Detailed observation of the cysts led us to classify them into three groups according to their morphology, structural integrity, and degree of immune infiltration: i) healthy-looking cysts with a rounded morphology and CD45 cells nearby; ii) cysts that started losing their characteristic morphology and displayed infiltration of immune cells; and iii) remnants of destroyed cyst free in the dermis (Fig.7b). We also observed that the proportion of cysts showing structural integrity markedly decreases with age, with a concomitant increase in the number of compromised or destroyed cysts, in which a prominent loss of epidermal markers such as E-cadherin is noted (red arrowheads) compared to rounded, “healthy” cysts (blue arrowheads) (Fig.7c). These observations were consistent with the aforementioned decrease in the total numbers of cysts in aged (18-month-old) mice (Fig.7c).

Of note, we observed that cells in cysts displayed abnormal nuclear morphology with characteristic DNA condensation patterns as well as positive staining for *γ*-H2AX (Fig.7d), indicative of DNA damage. In aged mice, where HFs are no longer synchronized, we detected cysts with irregular morphology showing positive staining for apoptotic markers (cleaved caspase 3 and TUNEL) (Fig.7d), indicative of the destruction and clearance of cysts from the GRK2 eKO aging skin via apoptosis at these latter stages.

Regarding CD45^+^ immune infiltrates, among the immune cell subpopulations present in cysts, we noticed a prominent presence of CD3^+^ T cells located both in the dermis surrounding cysts and within the epidermal cyst cell layers (Fig.7e), many of which were confirmed to be CD4^+^ T cells. Increased expression of adhesion molecules (ICAM1) in cysts might be mediating such immune cell recruitment. As reported in situations of HF immune privilege collapse [52], MHC-II-expressing immune cells are present in the vicinity of cysts, along with the presence of F4/80^+^ macrophages within destroyed cysts, surrounding cells, and wrapping their membrane around cell nuclei with irregular morphology (Fig.7e).

Collapse of the immune privilege and immune cell infiltration is one of the key features of the development of immune-mediated alopecias such as alopecia areata (AA) [53]. Strikingly, the phenotype of GRK2 eKO mice shared other several central hallmarks of AA. 18-month-old GRK2 eKO mice hair coat exhibited the presence of exclamation mark hairs (Fig.S8a), the prime diagnostic feature of AA [54], together with Keratin debris consistent with hair shaft fragility (Fig.S8b), or the presence of severe dermal inflammation involving polymorphonuclear cells such as neutrophils, macrophages and even foreign-body giant cells (black arrowhead, Fig.S8c). AA is also associated with leading melanin deposition in the dermis owing to the destruction of the follicular epithelium, termed as pigmentary incontinence [55]. Using electron microscopy and histochemical approaches we detected the presence of melanin in cysts at any stage, either inside healthy structures as well as free in the dermis or in the remnants of cysts that had already undergone immune-mediated destruction (Fig.S8d). Moreover, the skin of 8-month-old GRK2 eKO animals shared transcriptomic features of AA. The genes overexpressed in the skin of GRK2 eKO mice (GRK2 eKO signature) are significantly enriched in the affected skin with alopecia from the C3H mouse model, as well as in affected skin samples from AA patients (Fig.7f). Similarly, the genes overexpressed in affected skin samples from AA patients (human AA signature) are significantly enriched in the skin of GRK2 eKO mice (Fig.7g). Even when the GRK2 eKO phenotype shares several transcriptomic and histopathological similarities with AA, analysis of *GRK2* transcript levels in whole skin samples obtained from different types of AA patients (GSE68801) [56] did not correlate with disease (Fig.S8e). This could be due to the “dilution” of local GRK2 expression changes in whole skin samples, given its ubiquitous expression in different cell types, or reflect transient changes in different stages of disease progression.

Overall, our data indicated that GRK2 is essential for HF homeostasis, its deletion leading to complex alterations in HF cycling features and cell fate determination, eventually leading to the development of epidermal cysts that are destroyed during aging by mechanisms reminiscent of those operating in immune-mediated alopecias.

## Discussion

Tightly regulated spatiotemporal cell-cell and cell-niche intercommunications, as those involving Bulge-HG-DP crosstalk, take place via multiple stimuli and signaling cues that act through cell signaling networks and are responsible for maintaining normal HF homeostasis, cycling and cell fate determination [15, 17, 21, 22]. Using a keratinocyte-specific knockout mouse model, we uncover that GRK2 as a signaling node plays a central role in hair follicle biology.

A specific role for GRK2 during adult HF homeostasis and cycling is indicated by the facts that: i) the phenotype observed in GRK2 eKO mice is independent of HF morphogenesis; ii) cyst growth is only observed in cycling regions of adult GRK2 eKO animals, being absent in refractory and telogen-arrested regions as the ear skin [44]; iii) triggering global tamoxifen-inducible GRK2 ablation at the stage of second telogen in 2-month-old mice mimics the constitutive epidermal KO model.

Early alterations in GRK2 eKO HFs involve decreased hair germ size and unusual hair cycling features. Slow-cycling HFSC building up the bulge during telogen sharply differ from their activated counterparts residing in the HG [47]. GRK2 deletion selectively leads to HG shrinkage during the second telogen, leaving bulge HFSCs unaffected. Moreover, depilation experiments in p55 back skin indicated a delay in the telogen-anagen transition in GRK2 eKO animals, along with a retarded and heterogeneous progression in their proliferation pattern and anagen status, reduced HF length and pigmentation, all indicative of altered hair cycle progression, and consistent with the need of a fully functional HG to correctly achieve tissue regeneration.

HGs can be derived from catagen-surviving cells [4, 57] or from bulge cells migrating downwards during the catagen-telogen transition to acquire a HG fate [8, 58]. Although we observe equal or even slightly higher numbers of HFSC in GRK2 eKO mice at p55 compared to control, the possibility of a defective migration of quiescent SC to replenish HG cannot be ruled out. Since a smaller HG has been related with a delay in anagen entry [21], it may also be feasible that since we find that anagen entry takes place at later timepoints as a consequence of GRK2 deletion, HG formation during catagen-telogen transition might as well be delayed in GRK2 eKO mice.

Although both bulge and HG contribute to hair cycling during anagen, a smaller HG has been related with a delay in anagen entry [21]. The upper HG gives rise to ORS cells during anagen [59, 60], whereas the proximal part in close contact with the DP proliferates first by the end of the telogen phase and generates the matrix [15]. Matrix development is more severely altered than that of ORS in GRK2 eKO mice, suggesting that the region of the HG in closer contact with the DP would be more affected by the lack of GRK2. However, HFs with laser-ablated HG can recover normally with a minor SC input during anagen [14], suggesting that HG abnormalities alone will not account for our whole anagen delay phenotype, and that HFSC dysfunction due to the absence of GRK2 would play a role as well. Since HG may also contribute to keep the quiescent pool of stem cells in the bulge away from the DP-derived proliferative signals [47], a smaller HG may hamper such “isolation” and favor HFSC exhaustion and premature aging in the long-term [21], as observed in GRK2 eKO mice. Consistent with a central role for GRK2 within the stem cell niche compartment and during adult hair cycling, we have uncovered a dynamic modulation of GRK2 levels during the different hair cycle stages. A higher and more dynamic regulation of GRK2 transcript levels is noted towards the final phases of telogen in HG compared to bulge cells, and GRK2 protein levels are higher at 3 days post depilation in both HG and Bulge cells and remain high up to 8 days post depilation in advanced anagen IV-VI stages, suggesting that HG and anagen HFs are poised to upregulate GRK2 during telogen to anagen transition, while keeping GRK2 under a certain critical threshold would alter HG homeostasis, and thereby hair regeneration, as observed in GRK2 eKO mice. The signals and mechanisms underlying such dynamic changes in GRK2 expression remain to be investigated.

We postulate that the altered hair cycling features of GRK2 eKO animals give rise over time to the abnormal cyst-like structures, specifically emerging only in hair-cycling areas both by standard hair cycling as well as by depilation-induced hair regeneration. Despite originating from the lowest part of the HF (HG) and often remaining attached to the bulge (although not displaying HFSC labeling) GRK2 eKO cysts display aberrant growth and differentiation patterns. Cysts proliferate steadily in a hair cycle stage-independent way, are not destroyed in catagen, and display features of abortive HFs unable to fully acquire all the canonical anagen cell layers. Cysts can acquire ORS fate, one of the first structures produced during hair growth [3] and display as well markers of the matrix ascending companion layer, but the generation of IRS is compromised, probably linked to the absence of Gata3 expression, crucial for matrix cells to be transformed into the IRS layer [61]. Such defective IRS commitment may derive from an impairment in the generation of IRS progenitor structures or reflect a direct involvement of GRK2 in IRS maturation, both deriving into the formation of dystrophic hair follicles or cysts. Other authors [28, 37, 62–65] have reported contexts in which IRS is not correctly formed, typically within a phenotype affecting adult HFs but not hair morphogenesis. Why aberrant IRS formation debuts in the first’s postnatal hair cycles despite IRS fate is specified in a similar manner in both adult and morphogenesis stages [32] remains to be determined.

Our data also imply that although GRK2 ablation would lead to a general delay in hair cycle, as noted upon forced anagen induction through depilation, only a fraction of HFs would transform into cysts, whereas other follicles would undergo several rounds of hair cycles before developing cysts. Observing triple-bulged HF linked to cysts further strengthen this idea. Once the cysts start to develop, they proliferate steadily in a hair cycle stage-independent way, do not regress during catagen, and display an altered fate and lineage infidelity.

Strikingly, in GRK2 eKO mice, the DP rather than being engulfed by the HF remains located in contact with the proximal part of the dysplastic HF, being pushed away and displaced from the HG as cysts grow during anagen, thus promoting changes in HF stem cell architecture, linking cysts with hair loss. Our observations regarding cyst location always being beneath the newest bulge reinforces the idea of cyst emergence being associated with telogen arrest. However, contrary to what has been described in Hairless mice, GRK2 eKO animals show an evenly distributed proliferation status along cycle stages and cysts regions, in contrast to the proximal localization expected if this process was only dependent on paracrine DP stimuli. Therefore, it is tempting to suggest that the lack of cyst growth coordination with DP signals is determined by the lack of adequate responsiveness to DP-derived cues, rather than due to DP miss-localization as described [14, 66]. Overall, our data point to a self-generated growth potential within these epidermal cysts, independent from the growth-inducing cues emanating from the DP compartment.

In this regard, the fact that epidermal cysts arising in GRK2 eKO conditions display a multilayered epithelium expressing IFE-like differentiation and ORS/CL markers, but not IRS or late epidermal differentiation features, pointed to the occurrence of altered cell lineage determination processes. Klf5 and Sox9 have been described as master transcription factors (TF) demarcating IFE and HF lineages, respectively, and under homeostatic conditions its presence is mutually exclusive [18]. Strikingly, we find that both uni-lineage determinants are co-expressed inside cysts, a process known as lineage infidelity. The expression of Klf5 in cysts together with ΔNp63, which expression is typically restricted to IFE, may provide a causative explanation for their unresponsiveness to hair cycle-associated signals. Importantly, cyst cells not only display a shifted TF landscape, but also show a remodeled environment with the presence of tumoral-associated markers, and the stress TF c-Jun, which might contribute to the convergent expression of Sox9 and Klf5 [18].

It is tempting to suggest that cysts outgrowth in GRK2 eKO animals might phenocopy tumoral initiation stages, where lineage infidelity is known to provide cancer cell clusters with a heterogeneous and more plastic phenotype enabling tumor survival [46]. Bulge-targeted oncogene activation has been reported to drive tumoral growth specifically during anagen phase, when the environment is permissive with HFSC proliferation, concomitant with the emergence of cyst-like structures with marked similarities to our GRK2 eKO model [39]. The capacity of HF cells to give rise to tumors with mesenchymal features and higher malignant potential has been described [67]. In a previous study [43], we found that GRK2 loss in head and neck cancer cells triggers cell fate switching from epithelial to more mesenchymal features leading to tumor progression. Nevertheless, despite showing certain tumoral hallmarks, GRK2 eKO cysts do not further progress into SCC, at least in basal conditions, despite persistent lineage infidelity.

Our results put forward GRK2 as an integrative hub in signaling pathways related to HF homeostasis, in line with its reported participation in the modulation and crosstalk of multiple signaling networks in a cell type and context-dependent way [40, 42]. We suggest that GRK2 downregulation would simultaneously and cooperatively affect highly interconnected signaling cascades relevant to HF homeostasis and cycling, including decreased activation status of the Wnt/β-Catenin/Lef1 pathway, or unbalanced expression of transcripts related to networks involved in HFSC maintenance such as Wnt, BMP, or GPCR/adenylate cyclase/PKA pathways (see scheme in Fig.S9).

In this regard, truncated Lef1 expression in skin keratinocytes (ΔNLef1), which abolishes Wnt/βCat signaling [28], causes a mouse phenotype reminiscent to that of GRK2 eKO mice, with a delayed anagen, cyst growth and absence of IRS markers, consistent with the presence of Lef1 regulatory elements in hair-specific keratins promoters [68, 69]. Interestingly, GRK2 loss-of-function mutations in Jeune syndrome patients can lead to a defective response to Wnt ligands in chondrocytes due to the impaired phosphorylation of the Wnt co-receptor Lrp6 [70]. A BMP receptor KO mouse model (Bmpr1a KO) can correctly form the ORS but not the IRS layer during anagen, deriving in a multilayered epithelium full of keratin debris [34], with clear parallelisms with our mice phenotype. Spatiotemporally controlled Smad 2/3 and Smad 1/5/8 signals are required to achieve correct follicular homeostasis downstream TGFβ and BMP receptors, being the Smad2 cascade relevant during anagen induction [24] and the BMP signal required to form IRS [32]. Since GRK2 can modulate TGFβ signaling via its ability to phosphorylate Smad2 [71], and GRK2 downmodulation unbalances endothelial TFGβ networks [72], it is tempting to suggest that unbalanced Smads activation upon GRK2 loss may contribute to the observed hair follicle phenotype. Of note, labeling of ΔNp63, a signaling node related to Wnt and other pathways is detected in GRK2 eKO cysts, and overexpression of this factor prevents follicular cells to give rise to IRS and matrix cell populations and their shift in terms of epidermal and HF master regulators [73, 74]. On the other hand, GRK2 has been recently reported to exert a positive role in Hedgehog (Hh) signaling by relocating towards the ciliary base upon Smo GPCR-like receptor stimulation, allowing Smo phosphorylation and its subsequent interaction with the PKA-C catalytic subunit [75, 76], which eventually would relieve the inhibitory phosphorylation of GLI2, allowing the activation of downstream genes. Thus, GRK2 silencing would favor enhanced activation of the PKA cascade, consistent with the observed upregulation of GPCR/adenylate cyclase pathways signatures in our RNAseq data. It is worth noting that conditional epidermal deletion of Gαs (Gnas), a major regulator of the GPCR/Adenylyl cyclase/cAMP/PKA pathway, is alone sufficient to cause aberrant expansion of the stem cell compartment and rapid formation of basal cell carcinoma (BCC) in the skin, in a way dependent on decreased PKA activation status in such conditions, leading to activation of Hedgehog GLI and Hippo YAP1 transcriptional networks. On the contrary, persistent expression of the constitutive mutant active form GαsR201C in basal keratinocytes led with time to HF stem cell depletion and progressive hair loss via reduction of GLI and YAP1 transcriptional activity, with HFs terminally differentiating into keratinized cyst structures [77]. Thus, constitutively enhancing the adenylyl cyclase/cAMP/PKA pathway would favor the inhibitory phosphorylation by PKA towards the Hh cascade. Of note, the phenotype observed upon active Gαs expression shares common features with our GRK2 eKO model. Altogether, we propose that GRK2 downregulation could simultaneously affect the highly interconnected Wnt, BMP and Hh signalling pathways, thus impairing HF homeostasis.

Overall, the fact that several targeted genetic modifications involving crucial HFSC signaling components disturb hair follicles phenotype in ways closely resembling that triggered by GRK2 depletion, stresses the importance of GRK2 as a regulatory node in HF homeostasis and lineage specification during adult hair cycle.

Hair loss observed in GRK2 eKO animals starts much sooner than age-related hair loss [78], and appears to be linked to the progressive emergence of HF-derived cysts. Regarding the mechanisms involved, we uncover that GRK2 ablation in keratinocytes promotes key alterations in the HF stem cell niche. Mesenchymal-epithelial crosstalk enabled by the proximity between DP and HF is required for supporting epithelial cell growth throughout hair cycle [9, 12, 16, 79] and to maintain the molecular signature and hair-inductive properties at the stem cell niche [80]. In GRK2 eKO mice, the inability of cysts to respond to catagen-inducing signals leaves the DP in contact with the cysts but away from the HFSC niche, thus disturbing such homeostatic crosstalk. Given that the telogen HF relies in a close association between DP and HG to efficiently induce the entry into the growing phase, DP displacement would lead cyst-bearing HFs to become irreversibly telogen-arrested. Another possible additional mechanism would rely on the altered cyst fate, unable to normally signal back to the bulge promoting stem cell self-renewal and anagen entry, as happens when ORS and TACs are generated during the normal hair growth phase [21, 22].

These cumulative alterations triggered by cyst growth would have long-term consequences in bulge homeostasis and architecture. Changes in ECM properties and HFSC niche structural perturbations have been described to impact SC self-renewal in the context of aging, potentially connecting to HFSC loss [49, 81–83]. Compared to control mice, GRK2 eKO mice no longer show distinction between outer and inner bulge cell layers, with abnormal distribution of markers such as of E- and P-cadherins, ZO-1, β-catenin or F-actin, supporting the hypothesis of an aberrant organization of both cell layers affecting their functionality [84]. Such bulge structural aberrations progress from 8 to 18M, with unstructured HFSC clusters progressively lowering their CD34 levels, eventually leading to an unrecognizable bulge structure located below the sebaceous gland in aged mice. Thus, we postulated that the more hair cycles take place as mouse age, the more HFs would be transformed into cysts, hindering DP-bulge interactions and bulge functionality, and driving the hair loss phenotype in GRK2 eKO animals. Consistent with this notion, repetitive depilation exacerbates stem cell exhaustion and bulge destruction concomitant with macroscopic hair loss in GRK2 eKO mice.

Curiously, cysts emergence is not linked to development of spontaneous tumors with aging in GRK2 eKO mice, despite persistent lineage infidelity and several tumoral markers in basal conditions, reminiscent to that observed during wound healing or epithelial tumorigenesis [18], suggesting the existence of compensatory or counteracting processes. We find that GRK2 eKO cysts are destroyed during aging by mechanisms resembling those operating in immune-mediated alopecias.

The hair follicle bulge is known to be an immune-privileged site to avoid damaging long-lasting stem cells or its transient progeny [51, 53, 85]. The loss of such immune privilege is associated with different types of immune-mediated hair disorders, as alopecia areata (AA) or primary cicatricial alopecia (PCA), depending on the HF compartment displaying a collapsed privilege. Strikingly, key histological similarities are found between GRK2 eKO phenotype and AA, namely perifollicular CD4^+^ T-cells (“swarm of bees”), ICAM1 expression, the presence of dystrophic hairs (“exclamation point hairs”), or pigmentary incontinence. Similar to AA pathological mechanisms, bulge cells seem to be spared from immune-mediated destruction in our model. However, the main inflammatory targets differ between AA and GRK2 eKO mice, since the lymphocytic infiltration concentrates around HF-derived cysts rather than in the anagen HF proximal bulb.

Despite such differences in pathogenic processes, we have observed a significant correlation at the transcriptional level between whole skin GRK2 eKO RNA expression patterns and the ones present in AA patients during disease onset. Interestingly, cyst emergence can attract both intraepithelial and peripheral CD45^+^ cells even in the presence of an intact basal layer, owing to its intrinsic chemoattractant ability, which might be potentiated when immunogenic keratinocyte and melanocyte-derived peptides are released to the dermis once the basement membrane is discontinued during cyst destruction [52, 86], eventually leading to foreign body giant cell formation.

In the context of HF cells, lineage infidelity has been described to impact long-term hair homeostasis via triggering immune-mediated bulge cell destruction both in mouse models and human PCA follicular samples [87]. It is tempting to postulate that in our mouse model mixed fate cyst emergence triggers an immune-mediated attack, and that loss of immune privilege secondary to lineage infidelity might mechanistically explain AA proximal bulb apoptosis. Thus, our model points to long-term lineage infidelity and its association with stress-induced states as a major player in this process.

In sum, our work identifies keratinocyte GRK2 as a key player in HF homeostasis, cycling features and cell fate determination. Our data provide a better understanding on how abnormal HF-committed cells develop dystrophic follicular cysts and, on the mechanisms, linking cyst formation with hair loss, as well as suggest potentially interesting connections among these processes and the pathogenic mechanisms operating in immune-mediated alopecias. Further investigations are required to deepen in such connections and the underlying molecular mechanisms, and to explore the potential role of keratinocyte GRK2 in skin interfollicular homeostasis.

## Methods

### Experimental design

The objective of this study was to investigate the role of the GRK2 signaling hub in hair follicle biology. For this purpose, we have generated a mouse model with specific deletion of this kinase in keratinocytes and performed a thorough phenotypic and molecular characterization of these animals using a combination of histological, imaging and molecular biology approaches described in this section.

### Animal protocols

Mouse models used for these studies were housed and bred following all established regulatory standards, and all the experiments were performed in accordance to guidelines of the European Convention for the Protection of Vertebrate Animals used for Experimental and Other Scientific Purposes (Directive 86/609/EEC) and with the authorization of the Bioethical Committee of the Universidad Autónoma de Madrid and the Local Government (PROEX 339/14 and 118.8/21). The animal research facility of the CBMSO has the appropriate license to hold and use animals for scientific research (ES280790000180).

All the animals were bred and housed on a 12-hour light/dark cycle with free access to food and water. They were maintained at a room temperature of 22±2 °C with a relative humidity of 50±10% and under pathogen-free conditions.

#### GRK2^F/F^;K14-Cre mice

The C57BL/6 mouse strain carrying targeted GRK2 deletion in stratified epithelia was generated by crossing mouse expressing the Cre recombinase under the human Keratin 14 promoter (*K14-Cre*) [88, 89] with our mouse bearing GRK2 floxed alleles (GRK2^F/F^) [90]. The resulting hemizygous Cre positive males from this generation (F1) was then crossed to Cre negative hemizygous females to obtain a mixture of Cre positive and negative with floxed GRK2 in homozygosity (F2), which were then used for experiments and termed as GRK2 eKO.

<u>Tamoxifen-induced GRK2 knockout mice</u> were generated by crossing floxed GRK2 (*Grk2*^F/F^) mice with TxCre animals (from Dr. A. Kavelaars, University Medical Center Utrecht, Utrecht, the Netherlands), in which Cre recombinase expression is induced by the administration of tamoxifen. Floxed homozygous GRK2 mice (*Grk2^F/F^*) [72] and transgenic mice overexpressing a Cre recombinase fused to a mutant form of the estrogen receptor (ER) to allow for tamoxifen-dependent activation [B6.Cg-Tg(CAG-cre/Esr1*)5Amc/J] [91] were obtained from The Jackson Laboratory. *Cre^+/−^Grk2^F/wt^*offsprings were backcrossed with *GRK2^F/F^* mice to obtain mice carrying both floxed GRK2 alleles and, either none (*Cre^−/−^;Grk2^F/F^*) or a single CreER allele (*Cre^+/−^;Grk2^F/F^*).

For the experimental procedure, 2-month-old *Tx-CreGrk2^F/F^* mice were injected with 2 mg of tamoxifen (#T2859, Sigma Aldrich) in corn oil (#C8267, Sigma Aldrich) for 5 consecutive days to induce GRK2 depletion.

For <u>mouse genotyping</u>, genomic DNA extraction was performed by lysing a fragment of mouse tail in a lysis buffer containing EDTA (#E9884, Sigma Aldrich), SDS (CBMSO facility) and Proteinase K (#EO0491, Thermo Fisher) and using 2-propanol and ethanol (Merck) to precipitate the DNA. Then, mice were genotyped by using two different specific primers for PCR to detect either Cre (5’-CGATGCAACGAGTGATGAGGTTC-3’; 5’-GCACGTTCACCGGCATCAAC-3’) or GRK2 floxed alleles (5’-TGAGGCTCAGGGATACCTGTCAT-3’; 5’-GTTAGCTCAGGCCAACAAGCC-3’; 5’-CAGGCATTCCTGCTGGACTAG-3’). The products of amplification with the DNA dye as loading buffer 6X (#N472, EZVISION®) were subjected to electrophoresis on a 1-3.5% agarose gel in TAE and the resulting bands were detected using ultraviolet light.

### In vivo protocols

For proliferation analysis, BrdU (Roche) was injected intraperitoneally to mice at 0.1 mg/g weight in 0.9% NaCl 2 hours before sacrifice.

Recombination in *TxCre-Grk2^F/F^* mice was induced by an intraperitoneal injection of tamoxifen (Sigma-Aldrich, 2 mg per mouse, in 0.9% NaCl, containing 9% ethanol and 51% sunflower oil), for five consecutive days. For depilation experiments, synchronized back skin mouse was used. Hair was removed with an electric razor and then, Veet depilatory cream was applied for 1 min. For repetitive depilation experiments, this process was repeated every month, up to a total of 9 times.

### Histological procedures

#### Immunohistochemistry and immunofluorescence

After tissue harvesting, skin was fixed in 10% neutral buffered formalin (#HT501128, Sigma Aldrich) for 24h, then paraffin embedded and 5 µm thick sections were used for Hematoxylin (#7211, Thermo Scientific) and Eosin (#HT110332, Sigma Aldrich) staining (H&E), following conventional protocols described elsewhere. The rest of the tissue was embedded unfixed in OCT (Optimal cutting temperature) compound (#4583, Sakura) and frozen using dry ice. Immunohistochemistry of paraffin-embedded (FPPE) sections was performed following standard protocols with citrate buffer-mediated antigen retrieval and using biotin-conjugated secondary antibodies, Vectastain® ABC kit (#PK-4000, Vector laboratories) for peroxidase signal amplification and DAB (#SK-4100, Vector laboratories) for antigen visualization. Sections were then counterstained with hematoxylin and mounted in Sub-X mounting medium (#3801740, Leica).

#### Fontana-Masson melanin staining

For melanin staining, OCT tissue samples were sectioned at 12 µm thickness, with the use of a cryostat (#CM1950, Leica), and stained with the Fontana-Masson kit (#HT200-1KT, Sigma), following the manufacturer instructions.

#### Horizontal whole mount

For complete hair follicle visualization, immunofluorescence staining was performed on thick OCT sections (60-70 µm). Briefly, horizontal whole mount sections were fixed for 15 mins in 4% paraformaldehyde (#30525-89-4, Santa Cruz Biotechnology) in PBS, permeabilized for 10 minutes with 0.1% Triton X-100 (#X100, Sigma) in PBS, and blocked for 1 hour in PBS containing 0.05% Triton X-100, 1% Gelatin from cold water fish skin (#G7041, Sigma), 1,5% BSA (#3117332001, Roche) and 1,5% of donkey serum (#D9663, Sigma). Sections were incubated with primary antibodies (see table) overnight at 4°C, rinsed with PBS and incubated with secondary antibodies (see table) during 4h at room temperature, counterstained with DAPI (CBMSO microscopy facility) and mounted in Mowiol (CBMSO microscopy facility).

**Table 1.** List of antibodies used.

| Target | Source | Ref. | Use | Dilution |
| --- | --- | --- | --- | --- |
| GRK2 | Novus | NBP1-32534 | IHQ | 1/250 |
| GRK2 | Abcam | 137666 | WB | 1/1500 |
| β-Actin | Sigma | A5441 | WB | 1/1000 |
| α6 integrin/CD49f-conj | Invitrogen | 11-0495-82 | IF, FACS | 1/100 |
| Sox9 | Sigma | AB5535 | IF | 1/400 |
| Keratin 15 | Thermo Fisher | MS-1068 | IF | 1/300 |
| CD34 | Abcam | ab81289 | IF | 1/400 |
| P-cadherin | R&D Systems | AF761 | IF | 1/200 |
| Ki67 | Thermo Fisher | RM-9106-S | IF | 1/300 |
| CD34-eFluor (660) | Invitrogen | 50-0341-80 | IF, FACS | 1/100 |
| BrdU | Abcam | 6326 | IF | 1/250 |
| Lef1 | Cell Signaling | 2230 | IF | 1/100 |
| βCatenin | BD Biosciences | 610154 | IF | 1/200 |
| GRK2 | In house |  | IF | 1/400 |
| Lrig1 | R&D Systems | AF3688-SP | IF | 1/100 |
| Keratin 5 | Convance | PRB-160P | IF | 1/400 |
| Keratin 6 | Covance | 905701 | IF | 1/400 |
| CDP | Proteintech | 11733-1-AP | IF | 1/100 |
| Gata3 | Invitrogen | 14-9966-82 | IF | 1/100 |
| Klf5 | R&D Systems | AF3758-SP | IF | 1/50 |
| ΔNp63 | Biologend | Poly6190 | IF | 1/150 |
| cJun | Cell Signaling | 9165 | IF | 1/100 |
| Tenascin C | Sigma | T3413 | IF | 1/200 |
| αSMA | Dako | M0851 | IF | 1/50 |
| Gata6 | Cell Signaling | 5851 | IF | 1/250 |
| Filaggrin | Covance | PRB-417P | IF | 1/400 |
| Keratin 10 | Dako | M 7002 | IF | 1/100 |
| Loricrin | Covance | PRB-145P | IF | 1/400 |
| ZO-1 | Invitrogen | 40-2200 | IF | 1/100 |
| E-Cadherin | Cell Signaling | 3195 | IF | 1/300 |
| Faloidin Alexa Fluor 555 | Thermo Fisher | A34055 | IF | 1/100 |
| CD45 | Life Technologies | 14-0451 | IF | 1/250 |
| Y-H2AX | Cell Signaling | 9718 | IF | 1/150 |
| Cleaved Caspase 3 | Cell Signaling | 9661L | IF | 1/100 |
| CD3 | Biologend | 100201 | IF | 1/200 |
| ICAM1 | R&D Systems | AF796 | IHQ | 1/100 |
| MHC-II ( I-A/I-E) | Biologend | 107601 | IF | 1/200 |
| F4/80 | Abcam | 16911 | IF | 1/70 |
| Goat anti Rabbit IgG conjugated with Horseradish peroxidase | Nordicmubio | GAR/IgG(H+L)/PO | WB | 1/50000 |
| Goat anti Mouse IgG1 IgG2a IgG2b IgG3 conjugated with Horseradish peroxidase | Nordicmubio | GAM/IgG(H+L)/PO | WB | 1/50000 |
| Biotin-SP AffiniPure Donkey Anti-Goat IgG (H+L) | Jackson Immunoresearch | 705-065-147 | IHQ | 1/3000 |
| Biotin-SP AffiniPure Donkey Anti-Rabbit IgG (H+L) | Jackson Immunoresearch | 711-065-152 | IHQ | 1/3000 |
| Alexa 488 Donkey anti-Goat IgG | Thermo Fisher | A-11055 | IF | 1/250 |
| Alexa 488 Donkey anti-Mouse IgG | Thermo Fisher | A-21202 | IF | 1/250 |
| Alexa 488 Donkey anti-Rabbit IgG | Thermo Fisher | A-21206 | IF | 1/250 |
| Alexa 555 Donkey anti-Goat IgG | Thermo Fisher | A-21432 | IF | 1/250 |
| Alexa 555 Donkey anti-Mouse IgG | Thermo Fisher | A-31570 | IF | 1/250 |
| Alexa 555 Donkey anti-Rabbit IgG | Thermo Fisher | A-31572 | IF | 1/250 |
| Alexa 647 Donkey anti-Goat IgG | Thermo Fisher | A-21447 | IF | 1/250 |
| Alexa 647 Donkey anti-Mouse IgG | Thermo Fisher | A-31571 | IF | 1/250 |
| Alexa 647 Donkey anti-Rabbit IgG | Thermo Fisher | A-31573 | IF | 1/250 |

#### Proliferation assay

To detect BrdU incorporation the immunofluorescence protocol was modified, and permeabilization step was followed by 15 min incubation with HCl 2N at room temperature.

#### Lipid staining

For Nile Red staining, OCT sections (12 µm thick) were fixed for 5 min with paraformaldehyde, then incubated with Nile Red (#19123, Sigma Aldrich) at 2 µg/ml for 30 min, rinsed, counterstained with DAPI and visualized with the corresponding red fluorescence filter.

#### Apoptosis determination via TUNEL

Terminal deoxynucleotidyl transferase dUTP nick end labeling (TUNEL) (#12156792910, Roche) staining protocol was performed according to the manufacturer instructions. Briefly, 5 µm paraffin-embedded sections were deparaffinized, then the antigen retrieval was performed with citrate buffer pH 6, followed by incubation with the reaction mix with the label and enzyme solutions for 1 hour at 37°C and counterstained with DAPI.

### RNA sequencing and analysis

Total RNA from dorsal mouse skin was isolated from 5 control and 5 transgenic GRK2 eKO animals with RNAEasy Mini kit (#74104, Qiagen), following manufacturer instructions. Upon collection, skin was incubated overnight at 4°C with RNA Later solution (#10391085, Invitrogen) and then stored at −80°C. RNA-preserved tissue was thawed, and 20 mg were homogenized using metal beads with Tissue Lyser (Qiagen) in QIAzol solution (#79306, Qiagen). To avoid genomic DNA contamination, DNA digestion was performed in column with RNase-free DNaseI (#79254, Qiagen). RNA samples with RIN > 8 were used. RNA sequencing was performed after poly-A selection using Illumina NovaSeq 2×150 bp sequencing. Reads were aligned to GRCm39 mouse genome with “Rbowtie2”, counts extracted with featureCounts function from “Rsubread” and Gencode annotation release 27 (gencode.vM27.annotation.gtf). Raw and processed data are deposited at the GEO database under the accession GSE261821.

Differential expression between skin from control and GRK2 eKO animals was performed using DESeq2. We consider genes significantly over- or under-expressed if they had p-val<0.05 and adjusted pval<0.25. Gene Set Enrichment Analysis (https://www.gsea-msigdb.org/gsea/index.jsp) was used to analyze enrichment of genesets in transcriptome datasets. GSEA from RNA-seq was performed from DESeq2 normalized counts after removal of genes with counts=0 in all samples. Genesets were retrieved from MSigDB signature database (https://www.gsea-msigdb.org/gsea/msigdb/index.jsp) or were generated from in-house analyses. The “GRK2 eKO signature” used in Figure AA was constituted by 231 genes significantly overexpressed in the skin of GRK2 eKO mice when compared to Control animals, upon DESeq2 analysis (see above). The “Human AA signature” in Figure AA was composed of 353 genes overexpressed in AA samples (patchy type and transient patchy type samples) when compared with non-affected skin, upon Ttest analysis of the AA dataset GSE45513 retrieved from Gene Expression Omnibus (GEO) database (PMID: 27322477). Functional analyses were additionally performed with the ToppFun utility from the TopGene Suite (https://toppgene.cchmc.org/). Heatmap of individual genes was represented using MeV software (https://doi.org/10.1007/978-1-4419-5714-6_15). Graphical representation of selected GSEA was performed using ggplot2 in R [92].

### Data and materials availability

Raw and processed data are deposited at the GEO database under the accession GSE261821.

### Flow cytometry

For the preparation of a cell suspension of keratinocytes from the back skin, first, subcutaneous fat was removed using a scalpel. Afterwards, the epidermis was placed, side up, into a p100 dish with 0.25% trypsin solution (generated at our cell culture facility) and cultured overnight at 4°C. Epidermis was scraped and filtered through 70 and 40 µm cell strainers (Falcon) to obtain the single cell suspension by centrifugation. To sort the target cells, the cell suspension obtained was stained with CD34-eFluor660 and integrin a6 (CD49f)-FITC, for 1h at 4°C, in a buffer containing 2% FBS in PBS using a FACSAria Fusion (BSC II) device. Dead cells were excluded with the use of propidium iodide (#P4170, Sigma).

### Protein extract preparation and Western Blot

2-month-old mouse epidermal single cell suspension was pelleted and lysed in Laemli lysis buffer and protease (62.5 mM Tris base pH 7.4, 2% SDS (v/v), 10% glicerol (v/v)) and phosphatase inhibitor cocktail (#87785, Thermo Scientific).

Protein samples were mixed with 4X loading buffer, boiled for 5 min at 95°C and resolved by SDS-PAGE in 4–15% Mini-PROTEAN® TGX™ Precast Protein Gels (#4561086, BioRad). After electrophoresis, proteins were transferred to 0.45 µm nitrocellulose membranes (#1620115, BioRad) and then stained with Ponceau S (#141194, Sigma Aldrich). Membranes were blocked using 5% BSA (#3117332001, Roche) in TBS pH 8 buffer solution, and incubated overnight with primary antibodies (see table). Membranes were then washed with 0.1% Tween-20 TBS solution (#8221840050, Sigma Aldrich) and incubated with HRP-conjugated secondary antibody at room temperature for 1h. Membrane-bound secondary antibody was detected using Western Lightning ECL Plus (#NEL103E001EA, Perkin Elmer) and Medical X-Ray blue films (#FA158, Agfa). The bands were quantified by laser densitometry with a Biorad GS-700 scanner.

### Electron microscopy

Skin samples were fixed with 4% paraformaldehyde (Santa Cruz Biotechnology) and 2% glutaraldehyde (Sigma) in 0.1M phosphate buffer pH 7.4, for 2 h at room temperature and overnight at 4° C. Samples were rinsed with PBS and postfixed with 1% osmium tetroxide (Sigma) and 0.8% Potassium Ferricianyde (Sigma) in bidistilled water, at 4 °C for 1 h, then incubated with 0.15% tanic (Sigma) acid for 1 min at room temperature. After several washes, skins were incubated with 2% uranyl acetate (Sigma), for 1 h at room temperature, and dehydrated in a graded series of ethanol (Merck).

Infiltration of the resin was accomplished by incubating the samples in a mixture of ethanol and propylene oxide (#540021, Sigma Aldrich) 1:1 (V:V), for 5 min at room temperature followed by incubations in propylene oxide 10 min at RT. Then, the samples were incubated, for 45 min at RT, in a mixture (1:1; V:V) of propylene oxide and epoxi resin (#TAAB 812, TAAB Laboratories). Finally, the samples were incubated 1hr, at room temperature and overnight, in pure resin under agitation, and the next day incubated for 3h into renewed resin and placed into flat embedding molds. Polymerization of infiltrated samples was done at 60°C for 2 days.

Ultrathin sections of 70-80 nm were cut with an ultramicrotome Ultracut E (Leica) and mounted on Formvar-carbon-coated Cu/Pd 100-mesh or slot grids. Next, they were stained with uranyl acetate and lead citrate. A JEM1400 Flash (Jeol) transmission electron microscope was used at 100 kV, with a Oneview 4K x4K CMOS camera (Gatan).

### Microscopy

Fluorescence images were acquired with Zeiss confocal microscopes (Inverted LSM 700, Inverted LSM 800 or Vertical LSM 900). For widefield color images vertical AxioImager M1 coupled with a camera DMC6200 (Leica) was used. Image acquisition and processing was performed with the use of Zen software (Carl Zeiss) and Fiji (ImageJ).

### Statistics

Data is represented as mean values ± SEM of the indicated number of mice in the figure legend. Statistical significance has been determined using GraphPad Prism 8 software by two-sided unpaired Student’s t-test, for one-to-one comparisons, and by one-way ANOVA followed by the Bonferroni’s test in the case of multiple mean comparisons. Statistical significance as well as n is specified in each figure legend

## Acknowledgments

We thank Prof. Fiona M. Watt (EMBL Heidelberg) for her guidance during Alejandro Asensiós training at King’s College London in the initial stages of this work. We thank Cristina Delgado and Estefanía Vázquez Oro for their helpful technical assistance, and Pilar Hernández, Angustias Page and Carmen Segrelles for their excellent assistance with the histological processing of the samples at CIEMAT histology facility. The help from CBMSO Animal Care, Flow Cytometry, Electron and Optical and Confocal Microscopy facilities is also acknowledged.

This work was supported by:

Agencia Estatal de Investigación of Spain (grant PID2020-117218RB-I00 to FM) (H2020-MSCA Program, Grant agreement 860229-ONCORNET 2.0 to FM)

CIBERCV-Instituto de Salud Carlos III, Spain (grant CB16/11/00278 to FM, co-funded with European FEDER contribution)

Fundación Ramón Areces (to FM and CR)

Instituto de Salud Carlos III (grant PI22_00966 to C.R., co-funded with European FEDER contribution) Programa de Actividades en Biomedicina de la Comunidad de Madrid-S2022/BMD-7209 - INTEGRAMUNE (to FM)

AA held a PhD Fellowship from FPI programme Agencia Estatal de Investigación (PRE2018-084418) We also acknowledge institutional support to the CBM from Fundación Ramón Areces.

## Author contributions

AA designed and performed experiments, prepared figures, analyzed data, wrote and revised manuscript; MSF, JPG performed some experiments, analyzed data and prepared figures; KLA and JMP helped with study concepts and revised manuscript, RGE performed functional genomic analysis, provided study concepts, wrote and revised manuscript; FM Jr, and CR provided study concept and design, interpreted experiments, supervised the study, wrote and revised manuscript and obtained funding.

