## Supplemental figures for "The G-Protein-Coupled Receptor Kinase 2 Orchestrates Hair Follicle Homeostasis"

Fig.S1

**a**

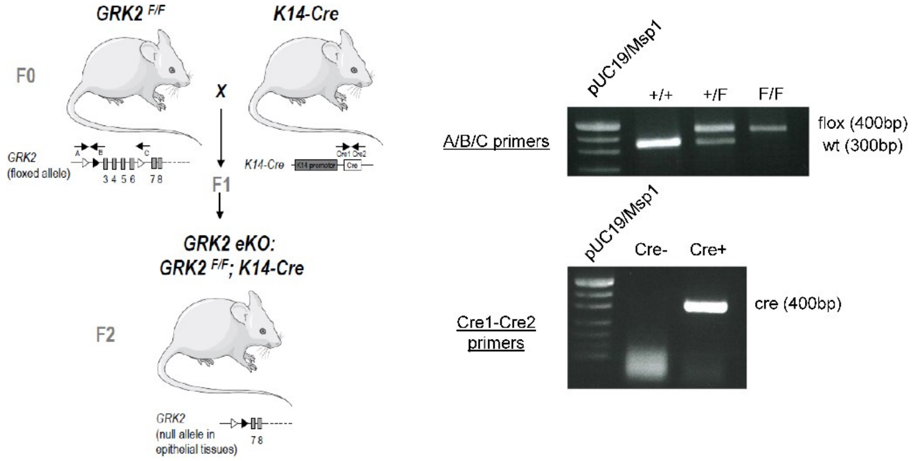
GRK2 F/F
K14-Cre
F0
X
F1
GRK2 eKO:
GRK2 F/F; K14-Cre
F2
GRK2 (floxed allele)
K14-Cre
GRK2 (null allele in epithelial tissues)
pUC19/Msp1
+/+
+/F
F/F
A/B/C primers
flox (400bp)
wt (300bp)
pUC19/Msp1
Cre-
Cre+
Cre1-Cre2 primers
cre (400bp)

**b**

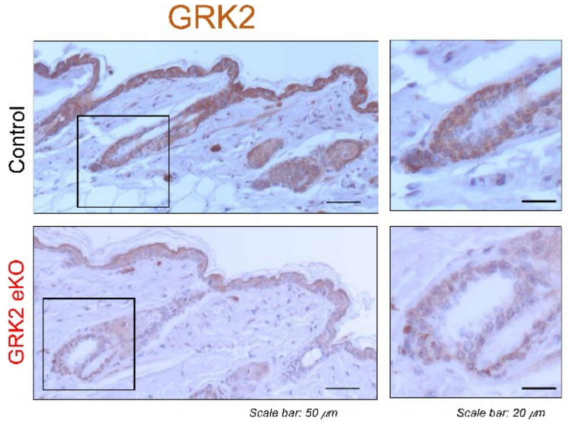
GRK2
Control
GRK2 eKO
Scale bar: 50 μm
Scale bar: 20 μm

**c**

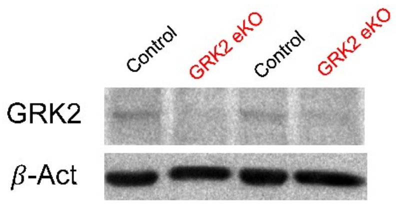
Control
GRK2 eKO
Control
GRK2 eKO
GRK2
β-Act

Fig.S2

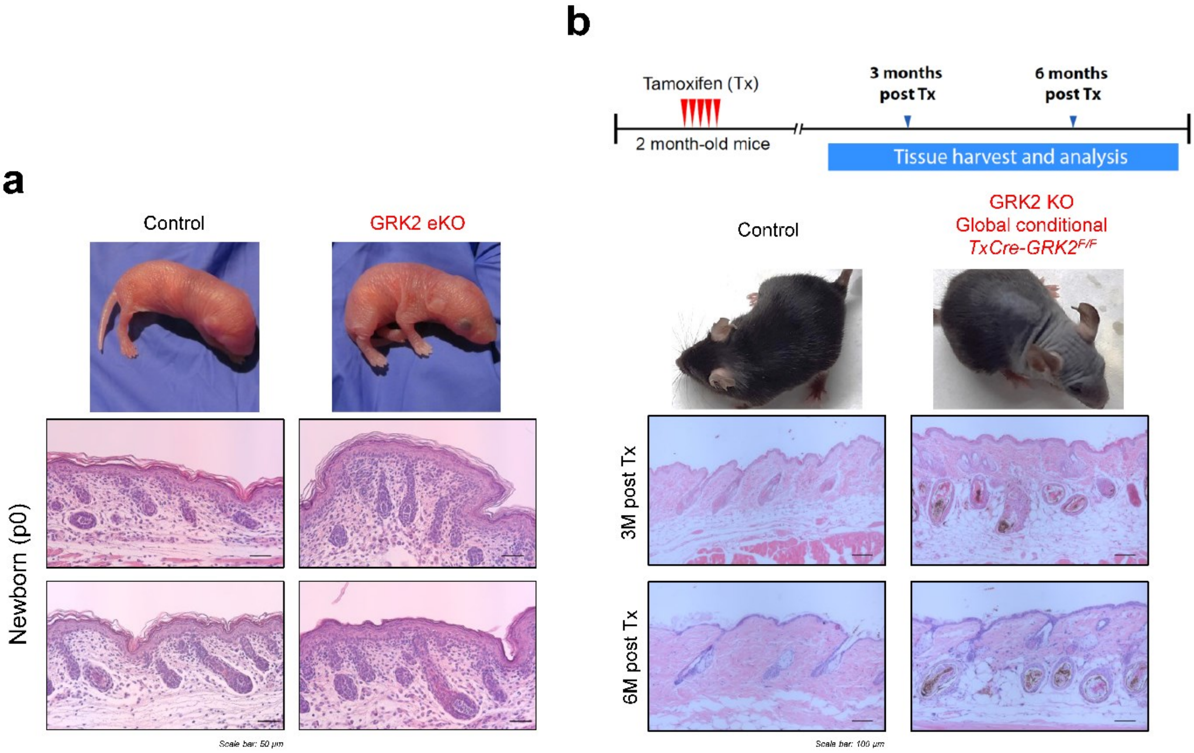
a
Control
GRK2 eKO
Newborn (p0)
Scale bar: 50 μm
b
Tamoxifen (Tx)
3 months post Tx
6 months post Tx
2 month-old mice
Tissue harvest and analysis
Control
GRK2 KO
Global conditional
TxCre-GRK2F/F
3M post Tx
6M post Tx
Scale bar: 100 μm

Fig.S3

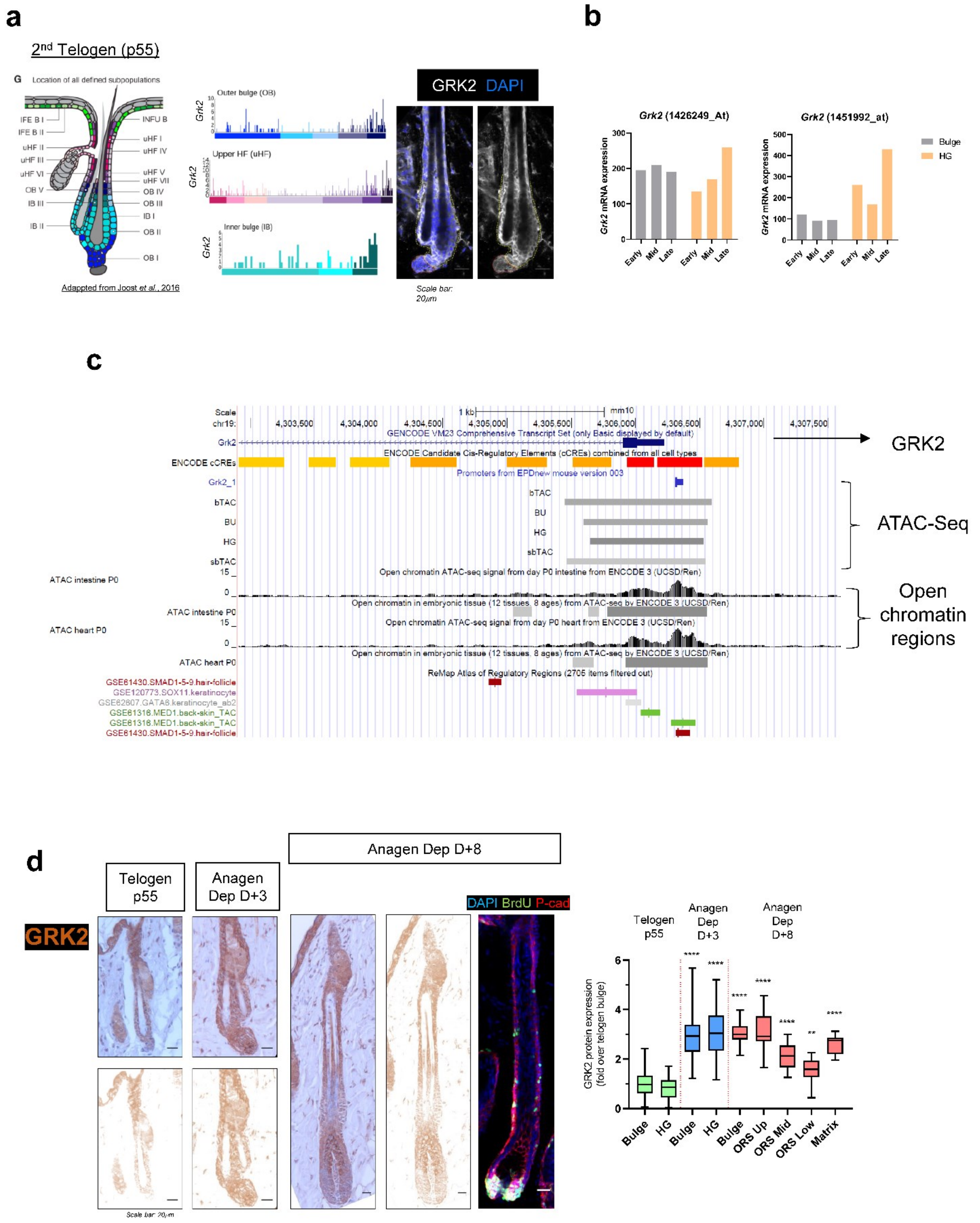

a
2nd Telogen (p55)
Location of all defined subpopulations
Adapted from Joost et al., 2016
Outer bulge (OB)
Upper HF (uHF)
Inner bulge (IB)
GRK2 DAPI
Scale bar: 20μm
b
Grk2 (1426249_At)
Grk2 (1451992_at)
Grk2 mRNA expression
Bulge
HG
Early
Mid
Late
c
GENCODE VM23 Comprehensive Transcript Set (only Basic displayed by default)
ENCODE Candidate Cis-Regulatory Elements (cCREs) combined from all cell types
Promoters from EPDnew mouse version 003
bTAC
BU
HG
sbTAC
Open chromatin ATAC-seq signal from day P0 intestine from ENCODE 3 (UCSD/Ren)
Open chromatin in embryonic tissue (12 tissues, 8 ages) from ATAC-seq by ENCODE 3 (UCSD/Ren)
Open chromatin ATAC-seq signal from day P0 heart from ENCODE 3 (UCSD/Ren)
ReMap Atlas of Regulatory Regions (2705 items filtered out)
ATAC intestine P0
ATAC heart P0
GSE61430.SMAD1-5-9.hair-follicle
GSE120773.SOX11.keratinocyte
GSE62807.GATA6.keratinocyte_ab2
GSE61316.MED1.back-skin_TAC
GSE61316.MED1.back-skin_TAC
GSE61430.SMAD1-5-9.hair-follicle
GRK2
ATAC-Seq
Open chromatin regions
d
Telogen p55
Anagen Dep D+3
Anagen Dep D+8
GRK2
DAPI BrdU P-cad
Scale bar: 20μm
GRK2 protein expression (fold over telogen bulge)
Bulge
HG
ORS Up
ORS Mid
ORS Low
Matrix

Fig.S4

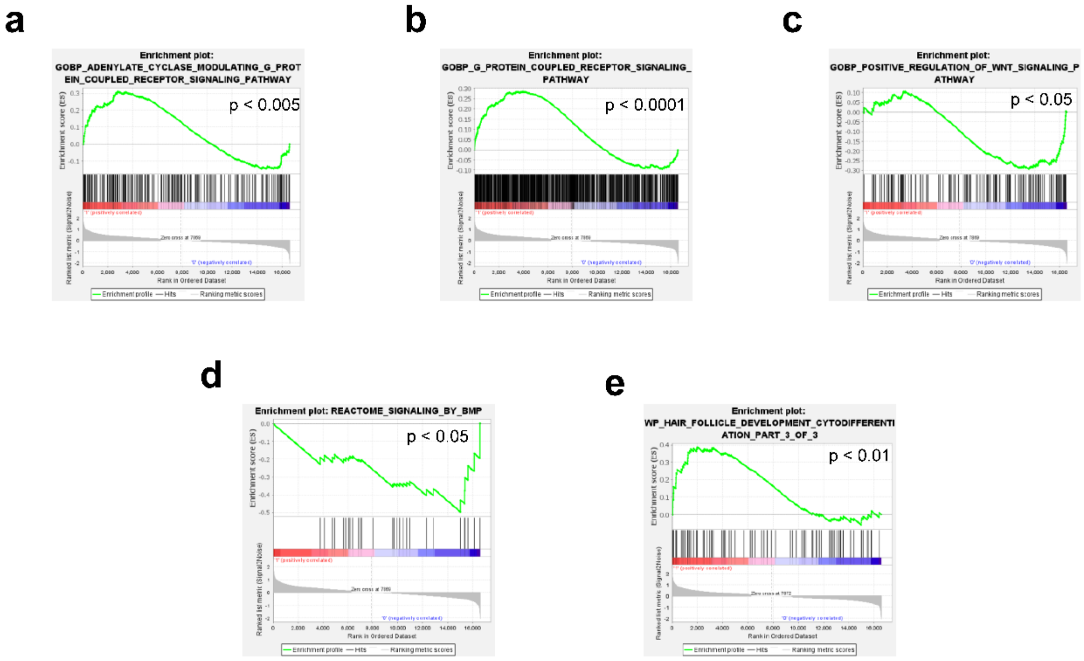
a
Enrichment plot: GOBP_ADENYLATE_CYCLASE_MODULATING_G_PROTEIN_COUPLED_RECEPTOR_SIGNALING_PATHWAY
p < 0.005
b
Enrichment plot: GOBP_G_PROTEIN_COUPLED_RECEPTOR_SIGNALING_PATHWAY
p < 0.0001
c
Enrichment plot: GOBP_POSITIVE_REGULATION_OF_WNT_SIGNALING_PATHWAY
p < 0.05
d
Enrichment plot: REACTOME_SIGNALING_BY_BMP
p < 0.05
e
Enrichment plot: WP_HAIR_FOLLICLE_DEVELOPMENT_CYTODIFFERENTIATION_PART_3_OF_3
p < 0.01
Enrichment score (ES)
Ranked list metric (Signal2Noise)
Rank in Ordered Dataset
Enrichment profile
Hits
Ranking metric scores

Fig.S5

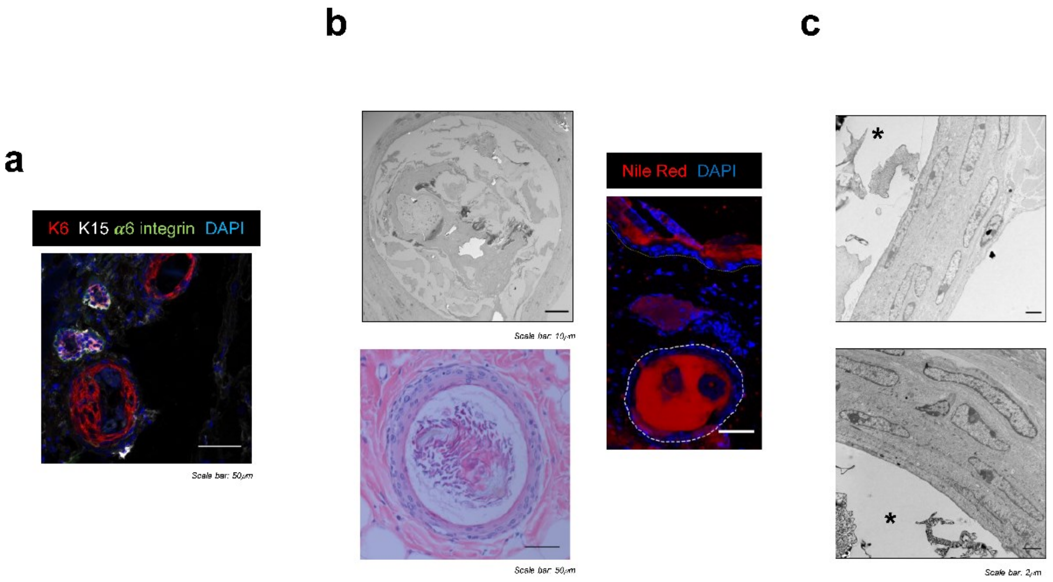
a
K6 K15 α6 integrin DAPI
Scale bar: 50μm
b
Scale bar: 10μm
Scale bar: 50μm
Nile Red DAPI
c
*
*
Scale bar: 2μm

Fig.S6

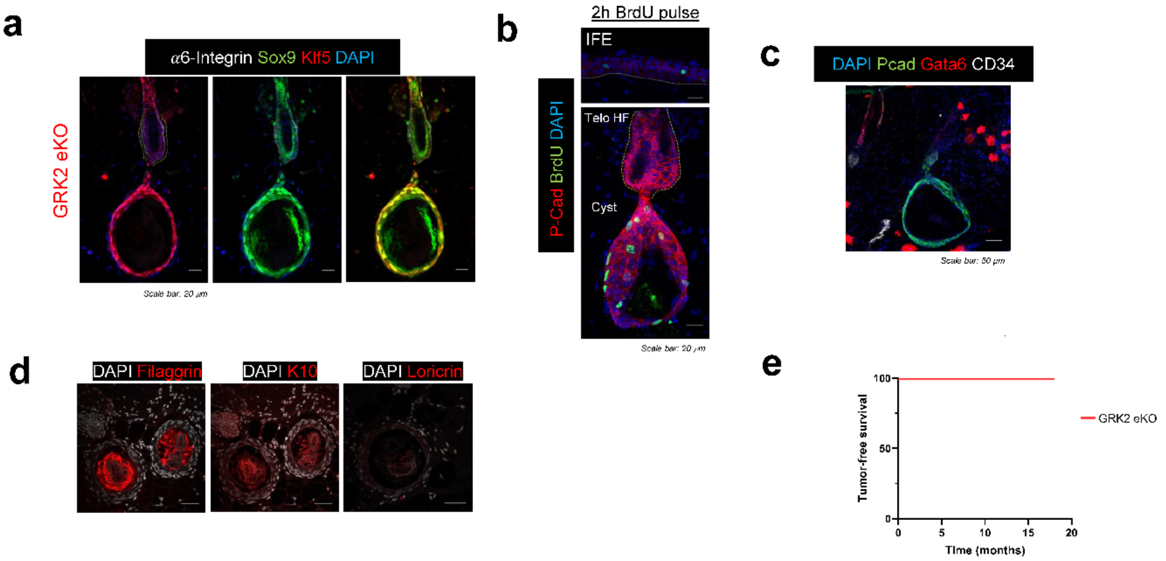
a
α6-Integrin Sox9 Klf5 DAPI
GRK2 eKO
Scale bar: 20 μm
b
2h BrdU pulse
IFE
Telo HF
Cyst
P-Cad BrdU DAPI
Scale bar: 20 μm
c
DAPI Pcad Gata6 CD34
Scale bar: 50 μm
d
DAPI Filaggrin
DAPI K10
DAPI Loricrin
e
Tumor-free survival
100
50
0
0
5
10
15
20
Time (months)
GRK2 eKO

Fig.S7

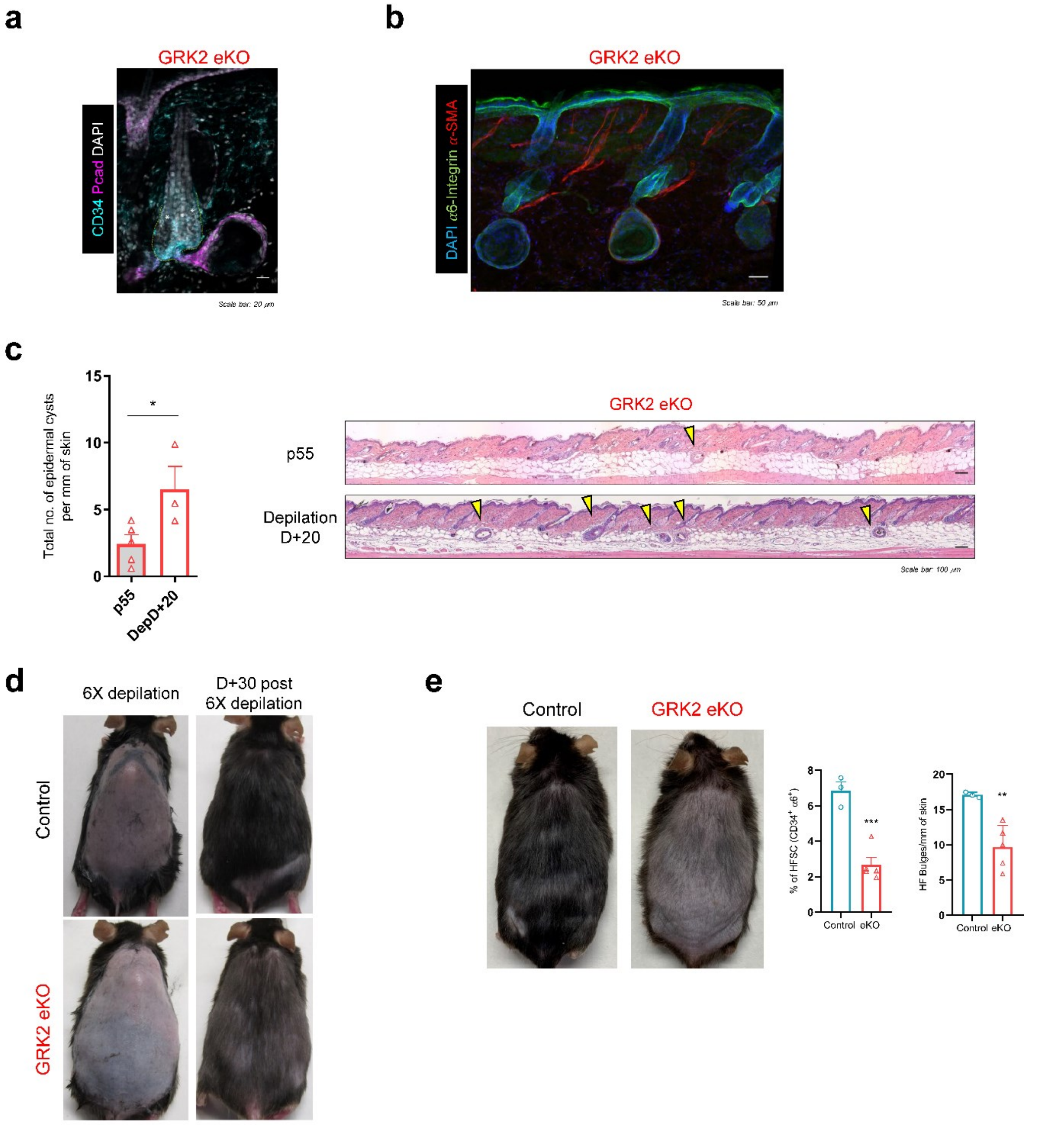
a
GRK2 eKO
CD34 Pcad DAPI
Scale bar: 20 μm
b
GRK2 eKO
DAPI α6-Integrin α-SMA
Scale bar: 50 μm
c
Total no. of epidermal cysts per mm of skin
15
10
5
0
*
p55
DepD+20
GRK2 eKO
p55
Depilation D+20
Scale bar: 100 μm
d
6X depilation
D+30 post 6X depilation
Control
GRK2 eKO
e
Control
GRK2 eKO
% of HFSC (CD34+ α6+)
8
6
4
2
0
***
Control
eKO
HF Bulges/mm of skin
20
15
10
5
0
**
Control
eKO

Fig.S8

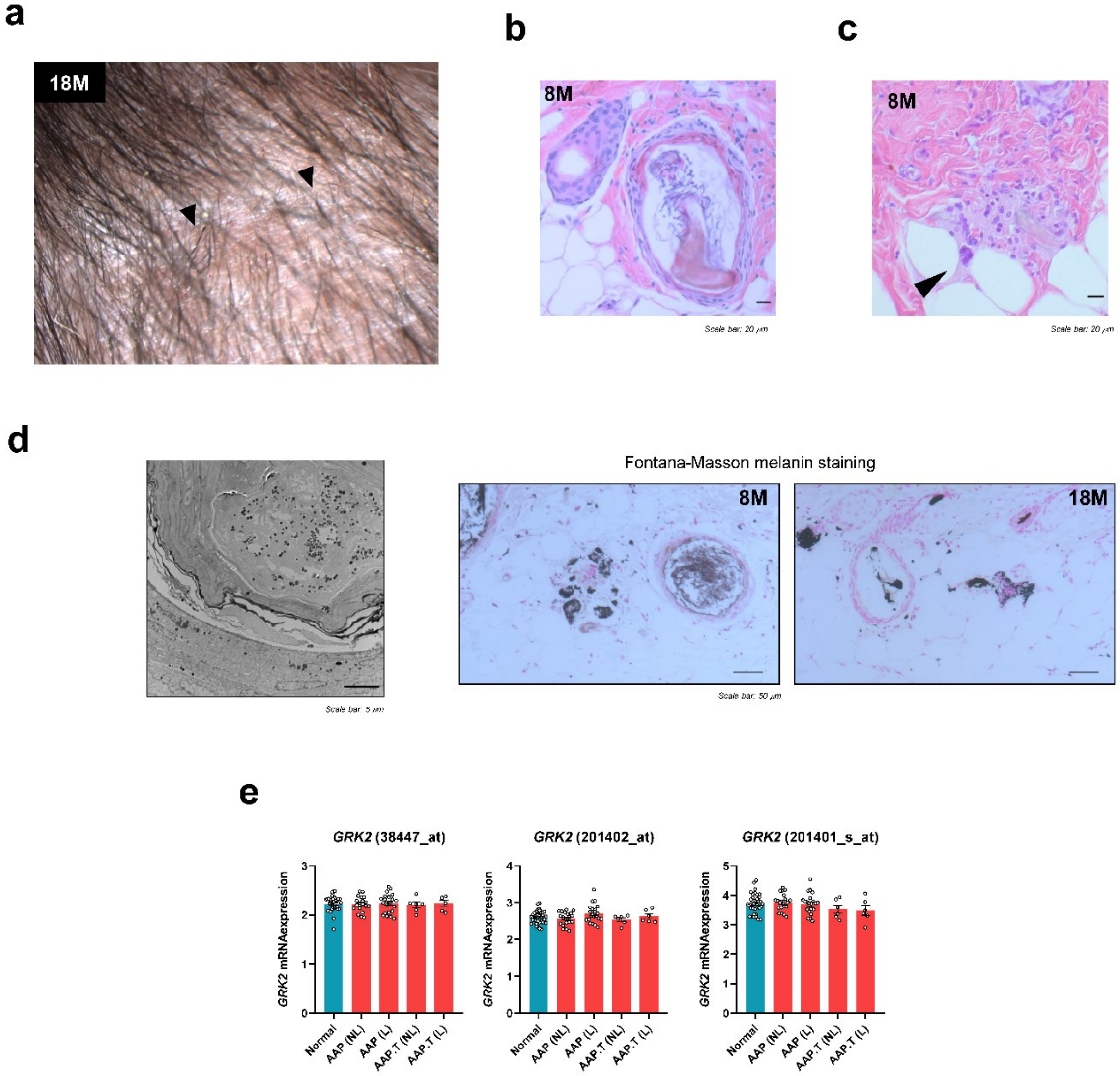
a
18M
b
8M
Scale bar: 20 μm
c
8M
Scale bar: 20 μm
d
Scale bar: 5 μm
Fontana-Masson melanin staining
8M
18M
Scale bar: 50 μm
e
GRK2 (38447_at)
GRK2 (201402_at)
GRK2 (201401_s_at)
GRK2 mRNAexpression
Normal
AAP (NL)
AAP (L)
AAP.T (NL)
AAP.T (L)

Fig.S9

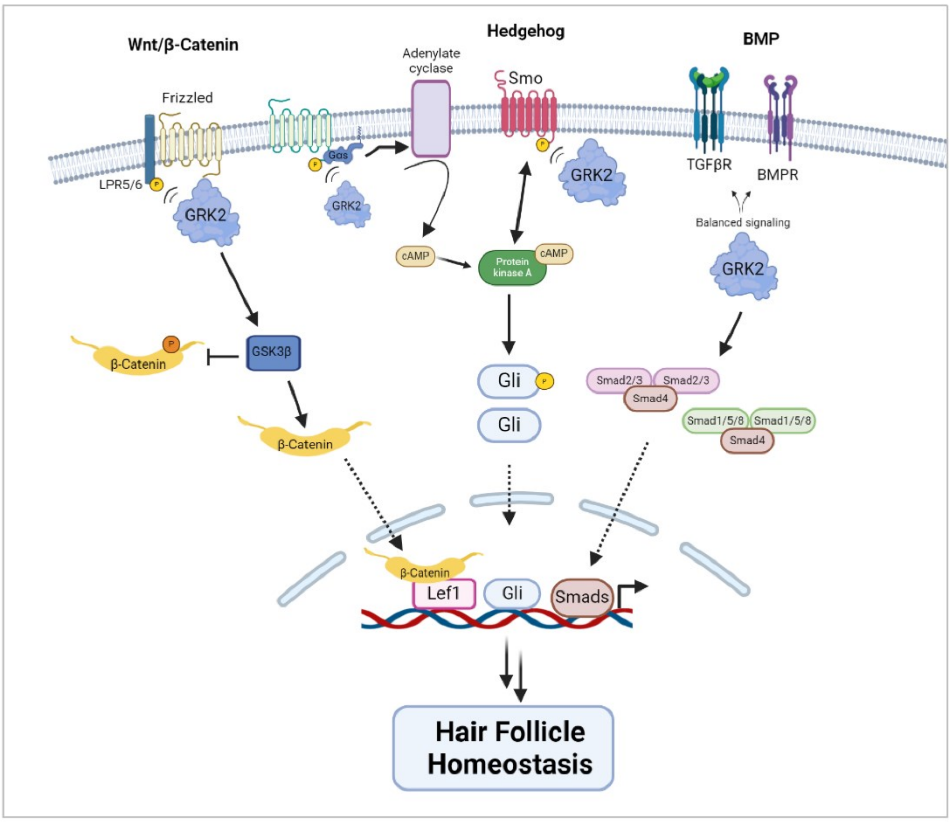

Wnt/β-Catenin
Hedgehog
BMP
Adenylate cyclase
Smo
Frizzled
LPR5/6
GRK2
Gαs
TGFβR
BMPR
Balanced signaling
cAMP
Protein kinase A
β-Catenin
GSK3β
Gli
Smad2/3
Smad4
Smad1/5/8
Lef1
Smads
Hair Follicle Homeostasis

### Supplementary figure legends

**Figure S1. Generation of an epidermal GRK2 knockout mouse model.** (a) Generation scheme of transgenic mouse model carrying GRK2 deletion in stratified epithelia and genomic characterization. (b) Confirmation of keratinocyte-specific GRK2 deletion via immunohistochemical detection of GRK2 in mouse skin sections or (c) by western blot of epidermal cell suspension.

**Figure S2. Observed hair loss phenotype is independent of hair follicle morphogenesis but dependent on postnatal GRK2 loss.** (a) Newborn (p0) GRK2 eKO pups show no alteration during embryonic hair follicle morphogenesis as assessed by histological analysis. (b) Schematic representation of Tamoxifen-induced global GRK2 deletion during second telogen (p60). This conditional model recapitulates epidermal-specific knockout hair phenotype suggesting a role for GRK2 during adult hair cycling.

**Figure S3. GRK2 levels in the hair follicle are physiologically increased during anagen.** (a) Resting hair follicles showed an enrichment of Adrbk1 gene expression in upper HF, upper bulge, and HG cells (database analysis of GSE67602 array), correlating with our immunofluorescence analysis for GRK2 protein staining at p55 in control animals. (b) Analysis of microarray data (GSE15185) of bulge and HG cells, where gene expression at different stages of the telogen phase (early, mid, late), revealed that HG cells display a dynamic Adrbk1 expression compared to the bulge throughout telogen, with higher expression levels at later stages. (c) A chromatin accessibility peak is found in primed HG cells compared to bulge stem cells in the GRK2 promoter (ATACseq data, GSE100876). (d) GRK2 protein levels increase during telogen to anagen transition in all HF populations compared to telogen phase bulge cells. DAB signal was quantified by image deconvolution using ImageJ to obtain the optical density of DAB channel. Means ± SEM data are represented, and statistical analysis was performed by 1-way ANOVA followed by Bonferroni's post-hoc test, $^{**}p < 0.01$, $^{****}p < 0.0001$.

**Figure S4. Wnt/β-Catenin, BMP, G-protein signaling, or Adenylate Cyclase pathways are deregulated in GRK2 eKO skin, leading to altered hair follicle differentiation.** (a-e) Significant enrichment in genesets related to BMP, Wnt/βCat, GPCR or adenylate cyclase and hair follicle cytodifferentiation in GRK2 eKO mouse skin.

**Figure S5. Hair follicle-derived cysts markers.** (a) Histological analysis of HF-derived cysts revealed their ORS and Companion Layer cell fate owing to the presence of Keratin 5 and Keratin 6 markers, together with the lack of HFSC marker Keratin 15. Note the orientation of sectioning was perpendicular to the HF and parallel to the interfollicular epidermis. (b) Cysts are also full of keratin debris and lipid secretions (Nile Red dye). (c) Cyst walls are comprised of a multilayered epithelium with elongated cell shapes and nuclear atypias as judged by TEM images.

**Figure S6 Cysts display mixed lineage markers and show an active IFE-like proliferation pattern but lack complete IFE differentiation features.** (a) Cysts display lineage infidelity, co-expressing a non-canonical mixture of IFE (Klf5) and HF (Sox9) markers, in contrast to bulges that express Sox9 alone. (b) Cysts show a constant proliferation pattern independent of hair cycle as judged by BrdU incorporation, further strengthening their mixed features resembling IFE. (c) Cysts also showed the absence of Gata6, marker of HF junctional zone. (d) IFE commitment has not been fully acquired as these structures remained negative for late differentiation markers

Filaggrin, Keratin 10 or Loricrin. (e) Kaplan-Meier diagram indicates the absence of spontaneous epidermal tumors in GRK2 eKO animals up to 20 months of age.

**Figure S7. Stem cell exhaustion and hair loss are coupled with hair cycles and aging in GRK2 eKO mice.** (a) Triple-bulged HF linked to cysts can be detected in GRK2 eKO animals, indicating that multiple rounds of hair growth are possible prior to eventual cyst emergence and its associated telogen arrest. (b) HFSC and DP physical distancing led to the presence of multiple cyst-bearing telogen-arrested HFs which are no longer able to begin the regenerative phase. (c) Analysis of cyst numbers found in p55 second telogen dorsal skin compared to 20 days post depilation (Depilation D+20) of GRK2 eKO mice. Quantification and representative images showed a significant increased in depilated conditions compared to telogen littermates. (d) Mice were subjected to repetitive depilation rounds. By the 6th depilation, GRK2 eKO mice lack the ability to replenish a complete hair coat, which appeared sparser at this point compared to control littermates. (e) After 9 consecutive rounds of depilation the inability to regrow their hair coat persisted in GRK2 eKO animals, which showed at this stage a significant reduction in HFSC number, as assessed by FACS-sorting CD34+ α6-integrin+ HFSC, as well as in the number of bulges, confirming stem cell exhaustion after several hair cycles. Means ± SEM data are represented, and statistical analysis was performed using unpaired t-test, $^{**}p < 0.01$, $^{***}p < 0.001$.

**Figure S8. Cyst immune-mediated destruction shares similarities with Alopecia Areata pathogenesis.** (a-d) Key Alopecia areata features are found along with destroyed cysts remnants in aged GRK2 eKO mice, such as exclamation dot hairs (arrows in a), keratin debris associated to hair shaft fragility (b), foreign body giant cells (arrow in c) and melanin granules free in the dermis as assessed by both electron microscopy and Fontana-Masson staining (d). (e) GRK2 transcript levels with different gen probes in skin punch biopsy samples obtained from (GSE68801). AAP (AA patchy type), AAP.T (AA transient patchy type), NL (non lesional), L (lesional). Means ± SEM data are represented, and statistical analysis was performed by 1-way ANOVA followed by Bonferroni's post-hoc test.

**Figure S9. Explanatory scheme.** Representative potential mechanistic connections linking keratinocyte GRK2 ablation with hair loss.
